# Marker direction alters genetic cell-state prioritization across complex traits: an endometriosis and chromatin case study

**DOI:** 10.64898/2026.11.13.766520

**Authors:** Elena Krikun, Abedalrhman Alkhateeb

**Author notes:** Corresponding author: Elena Krikun.

## Abstract

Genetic associations identify disease-linked genomic regions, but assigning these signals to cells depends partly on how marker genes are chosen. We compared genes selected by absolute fold change with genes up-regulated in each cell group and tested whether independent chromatin data supported the resulting interpretation. Reconstructing an endometriosis analysis, absolute-change markers prioritized venous endothelium, whereas up-regulated markers prioritized decidual stroma; the stromal result did not survive within-rule false-discovery correction. In a previously evaluated six-trait panel, neither marker rule consistently recovered the expected cell group. We then assessed nine additional genetic phenotypes with fixed marker definitions in one cross-tissue single-nucleus atlas. The leading cell annotation changed in four of nine phenotypes, but only two changes crossed broad cellular compartments; in one of those two, neither leader survived false-discovery correction. Complete cell rankings had correlations ranging from 0.523 to 0.843. Six broad cell-type proxies were recorded before this analysis; their median ranks were 24 with absolute-change markers and 20 with up-regulated markers, and their paired comparison was inconclusive (two-sided exact p = 0.125). Separately, chromatin accessibility in 36,073 nuclei from 12 endometrial biopsies supported stromal enrichment of inherited endometriosis risk in the initial analysis (z = 4.11; one-sided p = 0.00002). Later joint-model and biopsy-exclusion analyses supported the direction of that signal without identifying a unique causal cell state. Marker selection can change genetic cellular prioritization, but neither selection rule was universally superior.

**Teaser text:** The genes used to define a cell type can alter which cells appear linked to inherited disease risk. Across endometriosis and nine additional genetic phenotypes, marker direction changed cellular rankings, while independent chromatin evidence supported a broad stromal context in endometriosis.

## 1 Introduction

Genome-wide association studies (GWAS) reveal regions of the genome associated with disease. Turning these associations into a biological explanation is harder: a risk variant can affect a gene, but which cells are relevant? One common approach is to combine GWAS with a single-cell atlas and ask whether genes characteristic of a cell type carry more genetic signal than expected. Stratified LD score regression (S-LDSC) and MAGMA are frequently used for this step [1, 2]. Both depend on a decision that can look routine but has important consequences: how the marker genes are chosen.

A gene can be called a marker because it is expressed *more* strongly in one cell type, because its expression is particularly specific to that type, or simply because its fold change is large. The last definition, based on *|* log_2_ FC*|*, treats higher and lower expression in the same way. That can be useful for distinguishing cell populations, but it is a different biological question from asking which genes are active in a population. A set dominated by down-regulated genes can partly describe what a cell type lacks rather than what it expresses.

This distinction matters for endometriosis. The disease affects roughly one in ten women of reproductive age, yet the tissues and cellular processes through which inherited risk operates remain uncertain. Proposed mechanisms involve endometrial stroma and epithelium, perivascular cells, peritoneal mesothelium, and other developmental lineages. Large GWAS have identified loci near genes such as WNT4, GREB1, ESR1, and FSHB [3, 4]. Integrative studies have linked these associations to different cellular contexts: decidual and proliferative stroma, lesion macrophages, and SOX9-positive epithelium, depending in part on the dataset and analysis method [4, 5, 6, 7].

Recent work has compared statistical tools for linking GWAS to cell types and has developed ways to separate closely related cell populations [8, 9, 10]. Here we examine a complementary question: what happens *before* those tools run, when the marker set itself is defined? We use endometriosis as a detailed case study. We reconstruct a published cell-state ranking, separate the contribution of up- and down-regulated genes, and test seven marker definitions in a calibrated benchmark and a panel of known-source traits. We also compare marker-direction sensitivity across nine further genetic phenotypes in the same cross-tissue atlas, using a phenotype roster fixed before that analysis. We then examine genetic loci and ask how rankings change with donor sampling and across cell atlases. Finally, we test the broader stromal interpretation using chromatin accessibility from a separate group of donors. This last test selects accessible peaks without using RNA differential-expression marker sets, although marker genes are still used to label the ATAC cell clusters. Throughout, we distinguish the initial hypotheses from analyses added after earlier results were available. A related concern is that apparently reproducible computational signals may not identify a unique biological mechanism.

## 2 Methods

### GWAS

We used three endometriosis GWAS of European ancestry on GRCh37. GCST90205183 is the meta-analysis of Rahmioglu et al. without 23andMe [3].

KOLLER_EUR is the European meta-analysis of Koller et al. without 23andMe [4], downloaded from Zenodo (doi:10.5281/zenodo.18983492; CC BY-NC 4.0). FinnGen ENDO_ALL is the FinnGen R13 endpoint N14_ENDOMETRIOSIS (20,534 cases) [23]. Control traits were FinnGen R13 leiomyoma (CD2_BENIGN_LEIOMYOMA_UTERI; 42,816 cases), hypertension (I9_HYPTENS; 162,639 cases) and type 2 diabetes (T2D; 89,727 cases).

### Atlases and cohorts

We used HECA cells with counts [5]. The Koller cohort is the E-MTAB-14039 subset; the exact cohort is the library–individual pairs present in the HECA cell file with hormone-treated samples included (22/13). This matches the published sample counts; whether it is the identical set of experimental units cannot be confirmed from public metadata. The sensitivity cohort excludes hormone-treated samples. GTEx v9 snRNA-seq [24] was taken from CELLxGENE, keeping cell types with ≥50 cells and ≥3 donors. GSE266265 [12] (60 donors; donor × refined-type pseudobulk) is not part of HECA. GSE213216 [13] was read from raw 10x matrices.

### Differential expression

HECA and GTEx used scanpy [25] Wilcoxon tests, each type against the rest. In HECA this was done within disease status and compartment (endometrial vs immune), with ≤400 cells per type (fixed seed) and ≥30 cells. GSE266265 and GSE213216 used donor-or sample-paired pseudobulk: log2(CPM + 1) of the type vs the remaining cells of the same donor or sample, paired t-test, Benjamini– Hochberg correction within type. Eligibility was ≥10 donors with ≥10 cells (GSE266265) and ≥4 samples with ≥10 cells (GSE213216). For the HECA donor checks we also pooled labels over status with at most 50 cells per donor and type (donors with ≥10 cells; types with ≥3 donors), and ran donor-paired pseudobulk DE (moderated paired t with s0 = median standard error; ≥3 donors per status-split type, ≥4 donors per pooled type).

### Marker families

Seven families were built per type: ABS, *|* log_2_ FC*|* > 1.2 and adjusted p < 0.01 (the rule of Koller et al.); UP and DOWN, the same thresholds by sign; ABSN, UPN and DOWNN, the top 400 genes by absolute, highest and lowest score; and SPEC, each gene’s share of its CPM summed over types, as a continuous covariate conditioned on mean expression. Sets with fewer than 20 genes were dropped. The universe was genes mapped to a unique autosomal NCBI37.3 symbol.

### MAGMA

We used MAGMA v1.10 [2] with a 35 kb upstream/10 kb downstream window and the 1000 Genomes European reference. We ran competitive gene-set analysis conditioned on all tested genes, and gene-property analysis (direction = greater). Binary set indicators analysed as gene properties reproduced gene-set p-values (Spearman 0.97–0.98); they were used for conditioning and joint models (condition-hide). Trait contrasts compared standardized coefficients, one-sided:

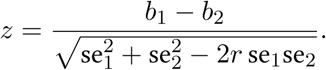

Here *r* is the correlation of the two traits’ coefficients over 1,000 paired circular genome shifts of the set (the same shifts for every trait). A contrast counted only if the empirical rotation p (share of shifted differences at least as large as the observed one) was also below 0.05. Gene-level covariates were log2 mean CPM, expression breadth (share of types with CPM ≥1) and gnomAD v2.1.1 LOEUF [11].

### S-LDSC

We used a Python 3 port of ldsc [1, 26] with baseline v1.2, 1000 Genomes Phase 3 European LD, HapMap3 SNPs, gene windows of ±100 kb, a 1 cM LD window and HapMap3 no-MHC weights. Each set was tested with --h2-cts against an all-tested-genes control. Joint fits added a competing set (focus, control, other). The sign decomposition fitted a type’s up and down parts jointly; a hit was down-carried if only DOWN|UP had p < 0.05, up-carried if only UP|DOWN had, and otherwise “both” or “neither”.

### Known-source benchmark

A planted profile (specific, depleted or highly expressed genes of a target state, scaled to the gene z-score SD) was added to real KOLLER_EUR gene z-scores. The null rotated the annotation layer (sets, covariates and profile) along genome order while keeping the real z-scores. We ran 300 null rotations and 40 replicates per alternative scenario (nine scenarios). Outcomes were top-1 exact state, top-3, correct group and true-state q < 0.05. FWER was reported with Clopper–Pearson intervals, recovery rates with Wilson intervals, and paired family differences with bootstrap intervals (10,000 resamples of replicates) and exact McNemar tests.

### Locus analyses

SNPs with p < 5 × 10*^−^*^8^ were clumped by distance (500 kb; GRCh37; autosomes). Candidate genes were MAGMA genes within ±250 kb of the lead SNP. Each locus was assigned to the compartment holding ≥90% of its expressed candidate’s specificity, else “broad”. Removal analyses dropped genes within ±500 kb of the removed loci; the null was 200 random locus subsets of the same size. Sharing with other traits was compared with 7 matched random windows per locus (pool of 800; matched on gene and SNP counts).

### Donor stability and dependence between GWAS

GSE266265 donor-paired DE was recomputed without each of the 60 donors. In HECA the ENDO-compartment Wilcoxon tests were rerun on the fixed cell sample with one endometriosis donor’s cells removed (no re-sampling), and the pooled designs left out each of 14 donors. Rankings used size-matched up-only sets and MAGMA. The effective number of donors in a group is 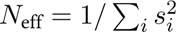, with *s_i_* the share of cells from donor *i*. The cross-atlas locus test reclassified the 57 loci with GSE266265 donor-mean CPM (same 90% rule) and removed the stromal loci of one atlas from the other atlas’s stromal set, against 200 random subsets of equal size. LDSC genetic correlations [26] used 1000 Genomes LD scores as reference and the HapMap3 no-MHC weights.

### Mesothelium selection (GSE213216)

Cell QC required ≥200 genes and < 25% mitochondrial reads. Mesothelial cells needed ≥3 of MSLN, UPK3B, CALB2, ITLN1 and PRG4, plus UPK3B or ITLN1, and none of PTPRC, PECAM1 or EPCAM. Other cells were assigned by the highest fixed-marker score; stroma was split by tissue context.

### Known-source panel

Four FinnGen R13 endpoints were added: I9_AF (67,783 cases), M13_DUPUTRYEN (6,972; official FinnGen R13 phenocode for Dupuytren contracture), C3_PROSTATE_WIDE (21,482; chosen by the registered name fallback because C3_PROSTATE_EXALLC was not in the R13 manifest) and E4_HYTHY_ AI_STRICT (56,098). Gene analysis was as above; p-values below 10*^−^*^300^ were set to 10*^−^*^300^ because MAGMA cannot read smaller values. Families and models were those of the GTEx v9 analysis. Expected groups were fixed before the run as regular expressions on cell-type labels, and the outcome was the rank of the best expected label. H-PANEL was supported if up-only markers ranked the expected group first in at least two thirds of the evaluable traits and more often than *|* log_2_ FC*|*; H-PANEL-N if size-matched up-only markers did so more often than size-matched *|* log_2_ FC*|*. Two label rules were corrected after the first run printed the labels, and both versions are reported.

### Additional cross-trait marker-direction analysis

The nine-endpoint FinnGen R13 phenotype roster and seven marker families were fixed before the expanded analysis; the secondary reference-proxy labels and descriptive metrics were recorded in a dated, immutable addendum before computation of cell ranks. Marker sets were constructed using the same GTEx v9 single-nucleus expression data and the corrected control-panel procedures, with the absolute-change, up-regulated, down-regulated, size-matched, and continuous specificity families unchanged. Gene-level MAGMA analyses used the existing European reference genotype panel and the build-37 gene annotation with 35-kb upstream and 10-kb downstream windows. Extremely small SNP *p*-values were floored at 10*^−^*^300^, as in the corrected earlier pipeline. Cell-type annotations were ranked by MAGMA gene-set or covariate association *p*-values, with Benjamini–Hochberg correction within each marker family. The DOWN family had 44 evaluable labels; the other six families had 45 each.

The primary descriptive measure was the proportion of nine phenotypes for which the leading label differed between UP and ABS. Prespecified descriptive secondary measures were Spearman rank correlation on the 45 shared labels, mutual ranks of the two first-ranked groups, and median and 90th-percentile absolute rank shifts. Six broad proxy labels were evaluated in a separate secondary comparison using the signed difference in natural logarithms of UP and ABS ranks. The two-sided sign-flip test enumerated all 2*^n^* assignments across nonzero paired differences, retaining ties for effect-size summaries. Five other rule-vs-ABS comparisons were adjusted using Holm’s procedure. We report results by both phenotype and prespecified overlapping phenotype group; neither unit should be interpreted as fully independent. The secondary proxy labels were locked before the completed expanded cell-ranking analyses. An earlier, aborted exploratory execution had downloaded the chronic obstructive pulmonary disease GWAS and started MAGMA gene analysis; its retained log stops at approximately 19.9% of processed genes, and the seven retained files from that aborted execution contain no completed gene analysis or cell-type rankings. Breast cancer appeared in its input configuration, but no breast-cancer execution is recorded. Thus, no earlier cell-ranking output is documented for either phenotype; this file audit cannot rule out unarchived outputs or unrecorded exposure. We therefore report the six-proxy comparison as exploratory, not an independent confirmatory test. The nine FinnGen phenotypes share participants and biological pathways, and the six overlap clusters do not provide independent effect-size replication.

#### Locus genes in lesions (GSE213216, GSE179640)

The lineage pseudobulk above (units with ≥10 cells) was used for eutopic endometrium, peritoneal lesions, endometriomas and unaffected peritoneum; samples of one donor and tissue were averaged. Each KOLLER_EUR locus gene (the expressed candidate used for its HECA compartment) was scored per sample as log2(CPM + 1) in its own lineage minus log2(CPM + 1) in the rest of the sample. The set statistic was the mean over expressed genes (mean CPM ≥1; at least 3 genes and 4 donors) and was compared with 10,000 random gene sets matched on expression decile, with locus genes excluded (one-sided permutation p). The rule was registered before the run: supported if the stromal-locus set exceeded its null in both lesion types. GSE179640 [14] was processed with the same rules from its 31 single-cell samples (organoid and bulk RNA-seq samples excluded); tissue was assigned from GEO metadata by a registered rule, and donor-paired comparisons with lesion-adjacent peritoneum and the mesothelium ranking followed the GSE213216 procedures.

#### Non-European GWAS

The East Asian, African and admixed American summary statistics of Koller et al. were mapped to rsIDs and analysed with MAGMA (35/10 kb windows), using the per-SNP effective sample size (sum of study weights) supplied with each file. The MAGMA site was not reachable, so we used the 1000 Genomes phase 3 superpopulation PLINK panels of Zenodo doi:10.5281/zenodo.6614170 (1.84 million variants), with gene annotation from each panel. Marker families were those of the GSE266265 analysis. HNE-EAS (eF3 size-matched up-only p < 0.05) and H-NE-ORIGIN (the originally documented candidate-origin criterion) were registered for the East Asian GWAS before the run.

#### Single-cell chromatin (GSE183771)

We obtained filtered peak-by-barcode matrices for 13 Cell Ranger ATAC v1.2 libraries (GRCh38) from GEO [17]. Two technical libraries came from the same biopsy and were treated as a single unit, giving 12 biopsies. We retained nuclei with at least 500 peak counts and no more than five times the median count for their library. Peaks called in different libraries were merged into one interval universe. We used the UCSC hg38ToHg19 chain to map intervals to GRCh37, accepting a lift only when at least 95% of bases mapped within one chain, the chromosome stayed the same, and the resulting interval was no more than twice as long as the original.

To identify cell clusters, we used TF–IDF on the 25,000 most frequently accessible peaks, followed by 30 SVD components, removal of the depth-associated first component, Harmony integration by library [18], and *k*-means (*k* = 30). We assigned cluster labels from accessibility around a fixed panel of 46 marker genes (gene body plus 2 kb upstream). A compartment label (mesenchymal, epithelial, endothelial, immune) required a highest score of at least 0.5 and a margin of at least 0.25 over the next compartment. Mesenchymal clusters were then considered stromal, perivascular or unresolved; epithelial clusters were classified as epithelium, ciliated or unresolved using the same margin. Unresolved mesenchymal cells were excluded from the stromal group and its comparisons; epithelial subtypes were pooled.

We summed peak counts by biopsy and lineage, requiring at least 20 nuclei per biopsy-lineage group. For each eligible group, we computed log_2_(CPM + 1) and subtracted the mean of the other eligible lineages in that biopsy. We then ranked peaks by a moderated paired statistic with *s*_0_ equal to the median across-peak SD. The top 5, 10 and 20% of eligible peaks formed the lineage annotations. Eligibility required an autosomal interval that had lifted to GRCh37, at least 20 total counts, and width no greater than 5 kb (240,290 peaks). The complete eligible set was the ALL control.

Four checks were fixed before inspecting any GWAS: (i) compartment composition within 10 percentage points of published values with no more than 10% unassigned nuclei; (ii) at least 95% successful lift-over of autosomal peaks; (iii) at least six biopsies contributing stromal and epithelial nuclei; and (iv) stronger overlap of stromal peaks with cultured stromal-cell ATAC peaks (GSE246585, all time points pooled) than of epithelial, endothelial and immune peaks, with a one-sided Fisher test against epithelium.

#### Chromatin S-LDSC

SNPs of the 1000 Genomes European panel were annotated by peak overlap and LD scores computed as above (1 cM window). Each lineage annotation was tested with --h2-cts together with the all-peaks control and baseline v1.2. The contrast between stroma and epithelium was tested in one model by re-parameterisation: with the stroma annotation, the sum of the stroma and epithelium annotations and the control in the model, the coefficient of the stroma annotation equals τstroma −τepithelium, and its block-jackknife standard error gives the test. The verdict rules (CH-1 to CH-3; trait-specific only if neither hypertension nor type 2 diabetes reached p < 0.05; a stopping rule if CH-1 failed) were fixed before the run.

#### Post-result joint scATAC model and biopsy-exclusion sensitivity

After the registered chromatin tests, we evaluated a five-annotation joint model comprising all accessible peaks (ALL), and the top-10% specificity sets for stromal, epithelial, endothelial, and immune nuclei. SNP overlaps were constructed in GRCh37 and LD scores were computed using the frozen 1000 Genomes Phase 3 EUR reference and 1-cM window. We fitted stratified LD score regressions with baseline v1.2 and baselineLD v2.2, retaining the original GWAS summary statistics, HapMap3 SNP definitions, and regression weights. Tests of positive stromal coefficients and the stromal-minus-epithelial contrast were one-sided. The latter used the full coefficient covariance matrix from the same joint fit; the software-generated coefficient names and covariance order were checked explicitly. We report nominal *p* values for these post-result analyses and do not interpret the related backgrounds or GWAS as independent tests.

For leave-one-biopsy-out (LOO) sensitivity, the original 13 GSE183771 ATAC libraries were reconstructed from their saved peak-by-cell sparse matrices and cell labels, pooling the two technical runs of biopsy S19a_18189. The eligible peak universe (240,290 regions) and original cluster labels were fixed. Counts from *all* 293,258 original universe peaks were used as the denominator for each biopsy-by-lineage CPM; after log_2_(CPM+1) transformation, specificity was the within-biopsy difference from the mean of other eligible lineages. Peaks were ranked by the original moderated statistic 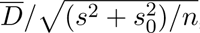, with *s*_0_ the median peakwise sample standard deviation and a stable top-10% selection. The full-data reanalysis was required to reproduce exactly the four frozen S10 peak sets, their chromosome-wise SNP masks, both M-vector definitions, and the primary KOLLER_EUR joint model. Each biopsy was then omitted as one block and all lineage-specific S10 masks and LD scores were recomputed, while the ALL mask, reference panel, GWAS, and cell labels remained fixed. The 12 LOO results were treated as correlated perturbations, not independent studies. The LOO procedure was added after the initial positive chromatin result.

#### Versioned analysis planning and deviations

Analysis choices, hypotheses, and verdict rules were documented in the versioned project analysis record before the corresponding runs, with earlier results available when some later plans were written. These are internal, run-specific records, not a publicly registered or externally time-stamped study protocol; a result obtained before a later plan was written did not become prospectively specified retroactively. Original confirmatory tests, later sequential follow-ups, audits, and analyses conducted after the first positive chromatin result are distinguished in the text and Table S1.

Hypotheses, falsifiers and outcomes are kept in a registry (Table S1). Deviations are recorded where they occurred: a mis-specified label rule, a cell cache lacking mesothelium, and two revisions of the chromatin cell labels made before any GWAS file was read. Results were appended to a results log after each run.

#### Human-participant data and ethics

This study used only existing, de-identified human genetic and single-cell datasets made available by the original investigators; no new specimens or participant-contact procedures were undertaken. Ethics and consent for the primary studies are described in their respective source publications.

#### Software

Python 3 (numpy, pandas, scipy, scanpy, scikit-learn, harmonypy, h5py, matplotlib), MAGMA v1.10 built from source on arm64, and the ldsc Python 3 port. All runs were on a single laptop with one worker.

## 3 Results

### 3.1 Marker selection changes the first-ranked cell type in a count-matched cohort

We began by rebuilding the study population used by Koller et al. The E-MTAB-14039 sample table contains 103 library–individual records spanning 63 libraries and 74 individuals; some libraries contain pooled samples. In the HECA cell file, 22 endometriosis and 13 control library–individual pairs met our matching rule when hormone-treated samples were included. These counts match those reported by Koller et al. (9 and 7 individuals, and 14 and 13 libraries). We use the term *count-matched cohort* deliberately: matching the numbers does not prove that every experimental unit or preprocessing choice was identical. To assess sensitivity to hormone treatment, we also analyzed the 18/13-pair subset that excluded treated samples.

We applied the published S-LDSC design to this cohort, testing 49 cell-state annotations under each marker-selection rule (Figure 1; Table 1). With genes chosen by the magnitude of their fold change, irrespective of direction, venous endothelium ranked first for KOLLER_EUR (Venous/ENDO: *p* = 1 *×* 10*^−^*^4^, *q* = 0.01); Venous/CTL and Arterial/CTL followed. Decidual stroma appeared lower in the ranking (dStromal_late/ENDO: *p* = 0.0067, *q* = 0.03). When we kept only up-regulated genes, dStromal_mid/ENDO moved to first place (*p* = 1.6 *×* 10*^−^*^3^), but its within-rule FDR-adjusted value was *q* = 0.08. With down-only markers, venous endothelium again came first (*p* = 9 *×* 10*^−^*^4^, *q* = 0.04). The key finding is a change in rank and interpretation, not a switch between two equally FDR-supported results. Disease-specific decidual states also had very uneven donor representation: one donor supplied 97% of dStromal_mid/ENDO cells (the donor-dependence analysis below).

**Figure 1:**
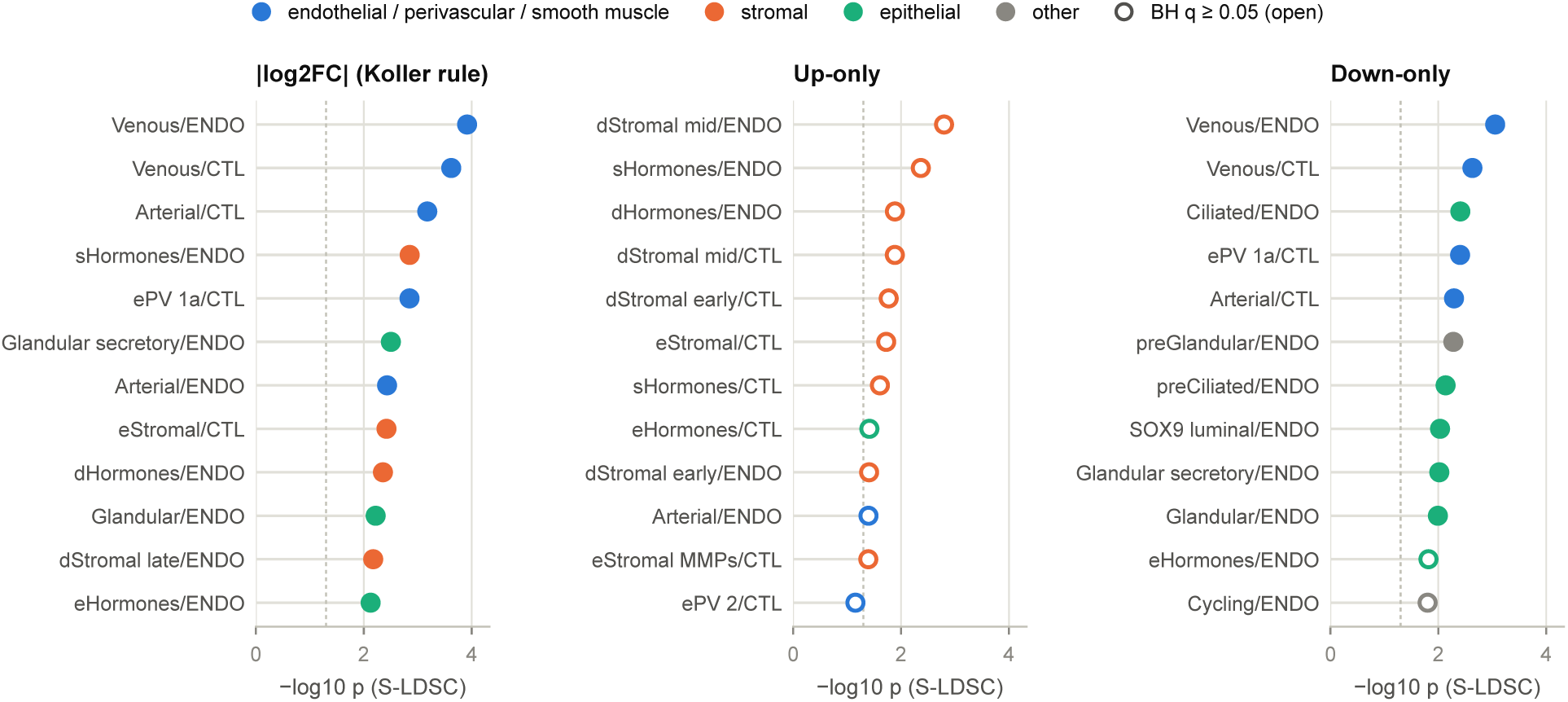
The first-ranked annotation flips with the marker-selection rule in the exact Koller cohort. S-LDSC with the pipeline of Koller et al. (KOLLER_EUR GWAS; 22 endometriosis and 13 control library–individual pairs, hormone-treated samples included). The 12 most significant annotations per rule; filled markers, BH q < 0.05 within the rule; dashed line, p = 0.05.

**Table 1:**
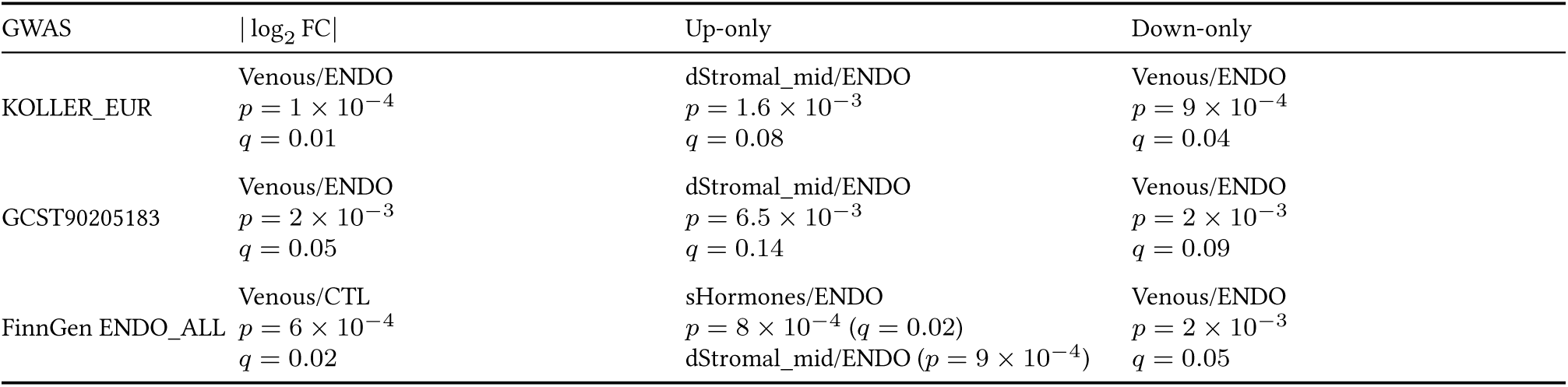
First-ranked annotation under each marker-selection rule in the count-matched Koller cohort. Within-rule BH *q* values are shown; the KOLLER_EUR up-only first-ranked state was nominal (*q* = 0.08). The experimental-unit identities of the reconstructed cohort could not be proven identical to those in the published analysis.

The difference persisted when we fitted competing cell-state annotations together (Figure 2). For *|* log_2_ FC*|* sets, no decidual or SOX9-functionalis set retained nominal significance (*p <* 0.05) against any of the six highest-ranked non-focus sets. In 33 of 36 eligible comparisons, only the competing set remained nominally significant; the competing sets survived every focus set. With up-only markers, both dStromal_mid/ENDO and dStromal_mid/CTL retained their associations against all six top competitors and against each of 11 vascular, perivascular, or smooth-muscle programs. dStromal_mid was also distinguishable from ePV_2. Excluding hormone-treated samples did not change the main pattern: the top *|* log_2_ FC*|* hit remained venous endothelium (*q* = 0.01–0.03 across tested cell variants).

**Figure 2:**
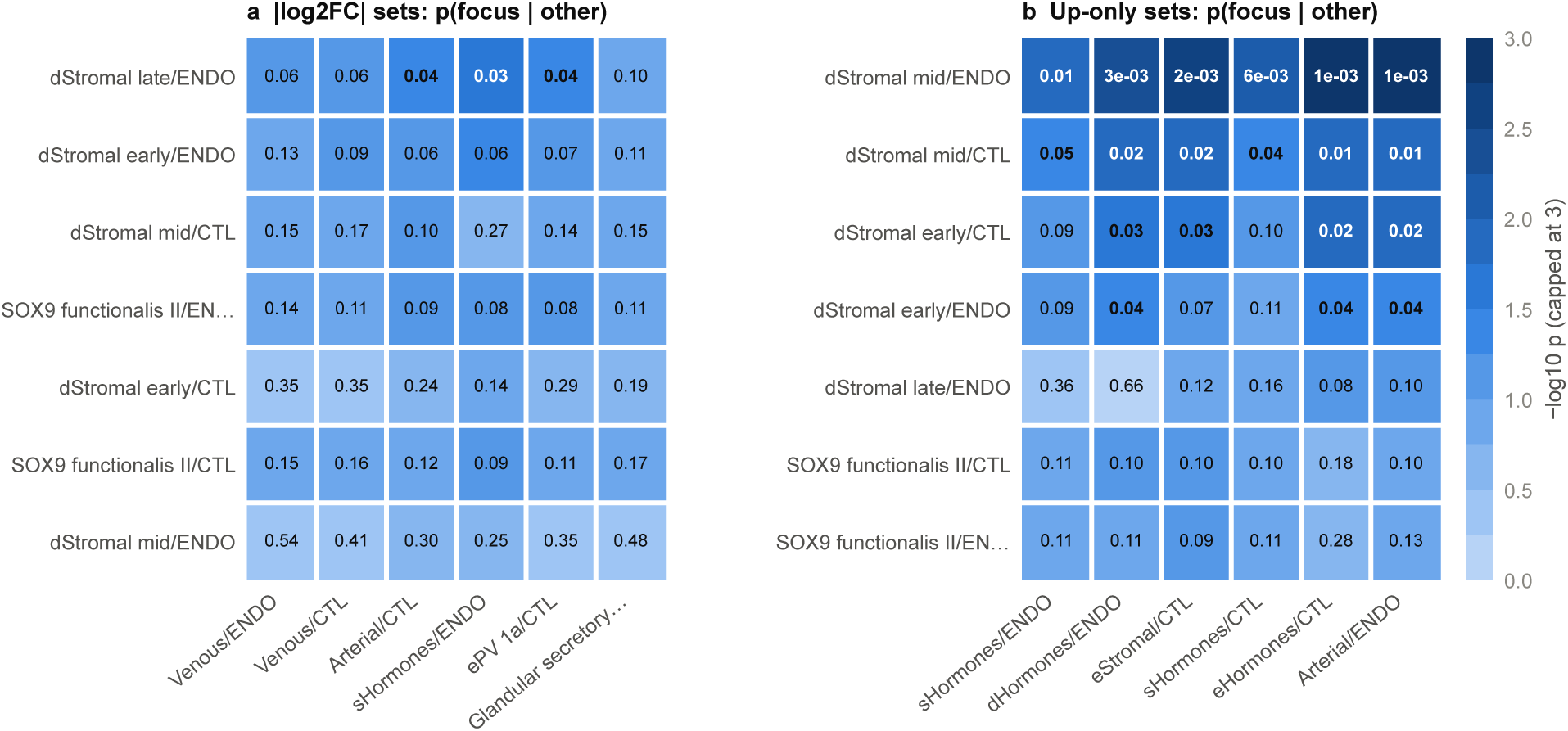
Joint models. Each decidual-stroma or SOX9-functionalis set (rows) fitted together with each of the six top-ranked non-focus sets of the same rule (columns), exact Koller cohort, KOLLER_EUR. Cell, p of the focus set given the other set (bold, p < 0.05).

### 3.2 Down-regulated markers can hide a signal in up-regulated genes

To understand the rank change, we split each bidirectional marker set into genes expressed more strongly and less strongly in the named cell type. In the count-matched cohort, the median *|* log_2_ FC*|* set contained 57% down-regulated genes. That proportion was 57–62% in the venous sets, 61% in Arterial/CTL, 36% in Arterial/ENDO, and 59–88% in the decidual sets. Among 34 nominally associated bidirectional annotations, joint S-LDSC attributed 19 primarily to their down-regulated portion, 8 to their up-regulated portion, 1 to both, and 6 to neither. The venous and arterial associations were carried mainly by the down-regulated genes; several early-decidual sets were carried by their up-regulated genes (Figure S1a–b).

This mixing could hide a biologically relevant signal. For dStromal_mid/ENDO, the complete *|* log_2_ FC*|* set was not nominally associated (*p* = 0.073), but its up-regulated portion was (*p* = 0.002). MAGMA showed a similar difference in HECA: the full set ranked 41st of 45 (*p* = 0.35; 83% of its genes were down-regulated), whereas the up-regulated portion reached *p* = 1.7 *×* 10*^−^*^4^. A positive-control example came from leiomyoma: the uterine smooth-muscle bidirectional set ranked second (*p* = 0.027; 60% down-regulated), but its up-regulated portion was much stronger (*p* = 4.7 *×* 10*^−^*^6^; Figure S1c).

The reverse process—a cell type appearing associated because of genes it expresses less strongly—was less general. The down-regulated portion was nominally associated in 27 of 30 top-five GTEx hits and 17 of 27 top-five HECA hits, but it was the sole carrier in only 10 of 30 and 10 of 27. This missed our prespecified threshold of at least one half. The composition of bidirectional sets also differed between atlases: GTEx sets contained a median of just 33% down-regulated genes, and their rankings more often tracked up-only than down-only rankings (six of six traits in HECA and five of six in GTEx). Even the same annotation could be classified differently by MAGMA and S-LDSC: Venous/CTL was down-carried in S-LDSC but carried by both directions in MAGMA. These findings support a concrete masking mechanism, not a claim that all bidirectional marker rankings are misleading.

### 3.3 A calibrated benchmark shows that no marker rule is best everywhere

A case study cannot tell us which rule will work in other settings. We therefore created nine benchmark scenarios by placing known cell-state signals into real KOLLER_EUR gene-level *z* scores. The sources were genes specific to, depleted in, or highly expressed in venous endothelium, decidual stroma, or perivascular ePV_2. We compared seven marker families: thresholded *|* log_2_ FC*|*, up-only and down-only sets; their size-matched versions (*N* = 400); and continuous expression specificity. To generate a realistic null, we rotated the complete annotation layer along genomic order while keeping the observed gene *z* scores fixed. An earlier attempt to shift the *z* scores instead had failed calibration and was not used. Across 300 null rotations, estimated family-wise error ranged from 0.3% to 1.7%; all Clopper–Pearson upper bounds were below 5% (for example, 3/300, 95% CI 0.2–2.9%). We used 40 replicates per alternative scenario, reported Wilson intervals, and made paired comparisons between methods (Figure S2; Supplementary Figure S3). With only 40 replicates, the maximum standard error of a recovery proportion is about 0.08.

- Specificity truths. Continuous specificity recovered the exact state most often (venous 0.93, ePV_2 0.78, decidual 0.47), then up-only (0.65, 0.50, 0.20); thresholded *|* log_2_ FC*|* managed 0.12, 0.12 and 0.03. The decidual rates carry wide intervals (specificity 0.47, 95% CI 0.33–0.63; up-only 0.20, 0.10–0.35). Up-only beat *|* log_2_ FC*|* for all three sources (paired differences +0.53, +0.38 and +0.17; McNemar p < 0.001, < 0.001 and 0.039).
- Depletion truths. Only down-only sets recovered the source (0.15–0.28); up-only and specificity families did not (0).
- Expression-level truths. No family exceeded 0.5 (best: specificity, 0.42).
- Size matching. It rescued much of the *|* log_2_ FC*|* deficit for venous endothelium (0.12 to 0.55) but not for decidual stroma (0.03) or ePV_2 (0.15). Size-matched up-only beat size-matched *|* log_2_ FC*|* clearly only for ePV_2 (+0.33, p = 0.002; decidual +0.12, p = 0.13; venous +0.05, p = 0.69).
- Related states. A decidual source was placed in the right stromal group in 82–90% of replicates but at the exact state in only 15–47%.

The thresholded *|* log_2_ FC*|* rule was never the best family in any of the nine scenarios. The direction of the true association decides which rule works; a rule that mixes directions is not the best in any scenario.

### 3.4 Positive controls reveal different strengths and weaknesses of marker selection

We next asked whether the choice of marker rule helps identify cell types that already have a strong biological connection to a trait. The controls were uterine leiomyoma, for which smooth muscle is expected, and hypertension, for which vascular and contractile cells are relevant. We defined the expected cell groups before running each analysis (Figure 3).

- Leiomyoma, HECA (MAGMA). Uterine smooth muscle ranked first with up-only, size-matched *|* log_2_ FC*|*, size-matched up-only and specificity sets, but not with thresholded *|* log_2_ FC*|* (rank 2).
- Leiomyoma, GTEx v9 (no uterus in the atlas). Smooth muscle ranked first with up-only, size-matched up-only and specificity sets, and 7th with *|* log_2_ FC*|*.
- Leiomyoma, exact cohort (S-LDSC). Uterine smooth muscle ranked 1st with up-only and 8th with *|* log_2_ FC*|*; nominal only (p = 0.011, q = 0.53).
- Hypertension. Vascular cells ranked first with every family except the down-only ones in HECA. In GTEx the prespecified pattern matched only specificity. Every other up-type rule ranked a “contractile cell” first, which a post hoc label check identified as pericytes and smooth muscle (author label Pericyte/SMC in 98% of nuclei); with that label all up-type rules and *|* log_2_ FC*|* passed.

**Figure 3:**
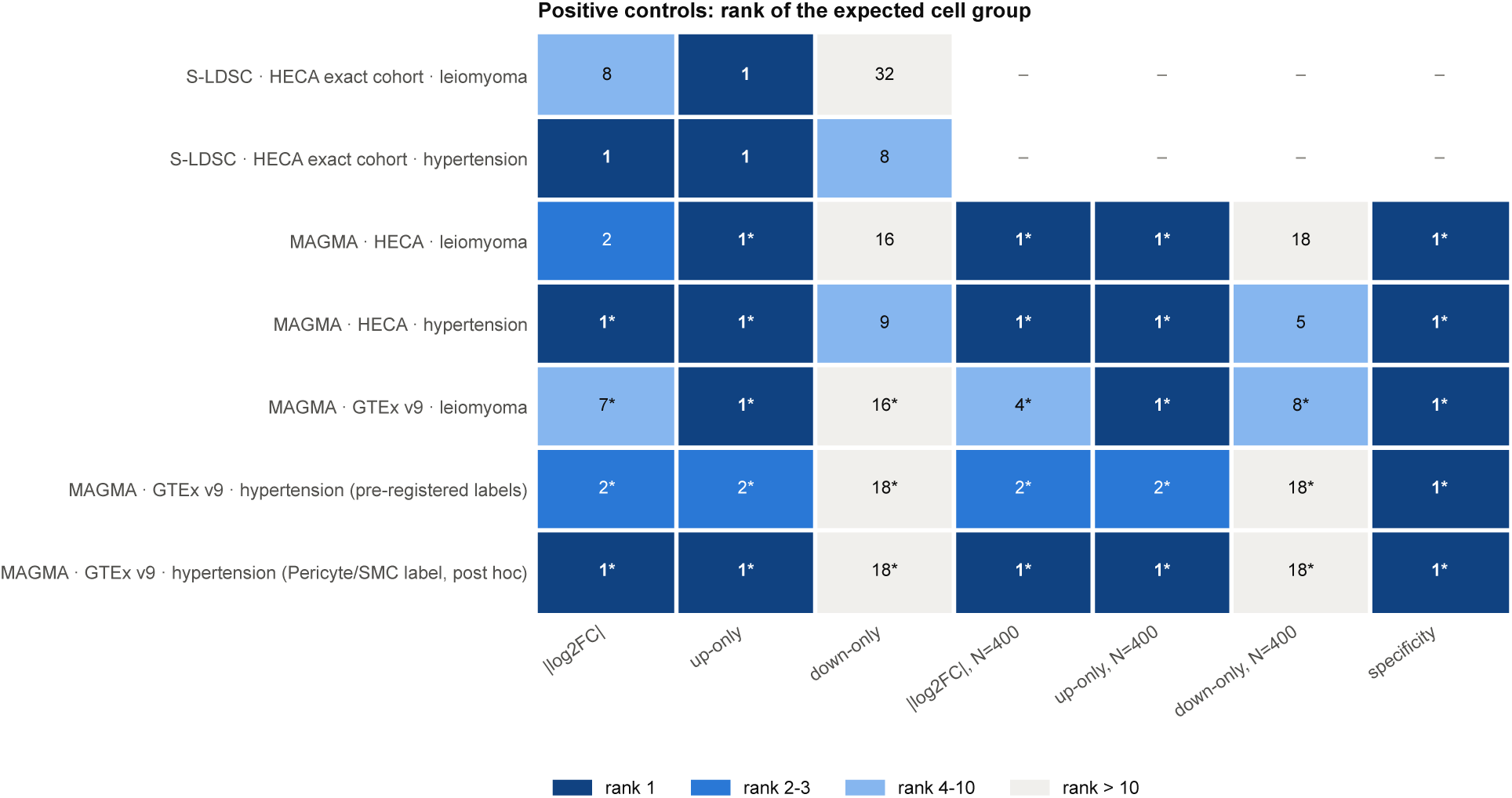
Positive controls. Rank of the best-ranked expected group (leiomyoma: uterine or visceral smooth muscle; hypertension: vascular, perivascular or smooth-muscle cells) per marker-selection family, in S-LDSC (exact cohort), MAGMA HECA and MAGMA GTEx v9. Asterisk, BH q < 0.05 within the family; dash, not run (S-LDSC was run for the three thresholded rules only). Expected groups were fixed before the runs; the last row uses the post hoc Pericyte/SMC label check. S-LDSC signals for the controls are nominal only.

#### Known-source panel

To test whether this generalizes, we added four FinnGen traits whose cell types were fixed before the run, all mapped in GTEx v9: atrial fibrillation (cardiomyocytes), Dupuytren contracture (fibroblasts), prostate cancer (prostate epithelium) and autoimmune hypothyroidism (lymphocytes) (Figure S4).

The results did not support a blanket preference for up-only markers. Across all six traits, thresholded *|* log_2_ FC*|* and up-only sets each placed the expected group first in three; the prespecified target for up-only markers was at least four and a clear advantage over the bidirectional rule. Size matching made a difference: size-matched up-only sets reached first place in four traits compared with two for size-matched bidirectional sets, and continuous specificity also succeeded in four. Failures differed by disease. The bidirectional rule missed fibroblasts in Dupuytren contracture (rank 15; this was the weakest GWAS and the best adjusted evidence was *q ≥* 0.10) and smooth muscle in leiomyoma (rank 7). Up-only markers placed skeletal muscle above cardiomyocytes in atrial fibrillation (cardiomyocytes ranked third), while the cardiomyocyte bidirectional set—only 6% down-regulated—ranked first. Prostate epithelium came first under both thresholded rules and their size-matched versions, and second under specificity. For autoimmune hypothyroidism, myeloid dendritic cells led under the thresholded rules and size-matched bidirectional sets; T cells led under size-matched up-only and specificity. We discovered two mis-specified cell-label matching rules after inspecting the list of 45 labels (lymphocyte matching inadvertently included mast cells, and the prostate rule missed the prostatic duct). The original-rule and corrected-rule outcomes are both retained.

#### Thresholds

Across eight threshold settings (*|* log_2_ FC*|* > 0.5, 1, 1.2 or 2; adjusted p < 0.05 or 0.01), up-only ranked uterine smooth muscle first for leiomyoma in 8 of 8; *|* log_2_ FC*|* did so in 2 of 8. For KOLLER_EUR, the first up-only label was a stromal state in 8 of 8 settings: decidual stroma in 5 and hormone-responsive sHormones stroma in 3. The prespecified target of decidual stroma in at least 6 of 8 was not met. At the most permissive threshold, the *|* log_2_ FC*|* rule ranked the same depleted-gene set (dStromal_early/CTL, 88% depleted) first for endometriosis, leiomyoma and hypertension.

#### Trait contrast

We also compared the strength of the decidual association with other diseases rather than just asking which annotation ranked first. The size-matched up-only decidual set had a larger standardized coefficient for KOLLER_EUR (0.036) than for type 2 diabetes (0.012) or hypertension (0.014). Covariance-aware one-sided tests gave *p* = 0.026 and *p* = 0.039; corresponding empirical rotation tests gave 0.006 and 0.015. The difference from leiomyoma (coefficient 0.020) was not significant (*p* = 0.081), consistent with shared uterine biology.

The contrast depended on marker definition. The bidirectional venous association was similarly strong for hypertension and endometriosis (0.032 and 0.035), whereas continuous decidual specificity was more strongly associated with type 2 diabetes than endometriosis (0.037 and 0.018). Contrasts in the smaller GCST90205183 GWAS were not significant. Covariance was estimated from 1,000 paired genomic rotations; cross-trait coefficient correlations were low (*r* = *−*0.01 to 0.14), so accounting for participant overlap made little difference to these particular *p* values (Supplementary Figure S5).

#### Generic gene properties

Conditioning on mean expression, expression breadth and gnomAD LOEUF [11] removed much of the significance in both atlases, but the first-ranked label of every rule stayed the same for KOLLER_EUR (Figure S1). The up-only and size-matched decidual sets kept p < 0.05 (1.2 × 10*^−^*^3^ and 8.9 × 10*^−^*^4^). The disagreement between rules is therefore not driven by gene-level confounders.

### 3.5 Marker-direction sensitivity extends across nine additional genetic traits

We next evaluated nine additional FinnGen R13 phenotypes in the same GTEx v9 single-nucleus atlas, using the previously defined seven marker rules and 45 cell-type annotations. The nine-phenotype roster was locked before this analysis; six broad cell-type proxies were additionally recorded in an immutable addendum before the new cell rankings were computed. The secondary proxy analysis remains exploratory rather than a claim of independently established causal cell types. The prespecified descriptive endpoint was whether the first-ranked cell-type annotation differed when marker genes were selected by positive fold change (UP) rather than absolute fold change (ABS).

The first-ranked annotation changed in four of nine phenotypes (44.4%): asthma (myeloid dendritic cell to T cell), breast cancer (fibroblast of cardiac tissue to bronchial epithelial cell), heart failure (slow muscle cell to macrophage), and varicose veins (cardiac endothelial cell to lymphatic endothelial cell). Two of the four were within broadly defined compartments (immune for asthma, endothelial for varicose veins). Thus, a shift between broad cellular compartments occurred in only two of nine phenotypes (breast cancer and heart failure); neither heart-failure leader survived within-family false-discovery correction (ABS *q* = 0.071, UP *q* = 0.161). The other five traits retained the same leading annotation, but this does not imply identical rankings. The nine Spearman correlations of the full 45-cell rankings ranged from 0.523 to 0.843 (median 0.737), and within-trait median absolute rank differences ranged from three to nine positions. Both leaders for non-small-cell lung cancer were also non-significant after adjustment (*q* = 0.333 and 0.148); its highest-ranked label was prostate luminal epithelium rather than a lung cell. The breast-cancer ABS leader was a cardiac fibroblast, likewise illustrating that a highly ranked label need not be anatomically plausible. The overlapping phenotypes do not provide independent replication (Table S2).

For the six broadly defined cell-type proxies fixed in the pre-analysis addendum, UP improved the proxy rank in four phenotypes, worsened it in one, and tied in one relative to ABS. The median proxy ranks were 24 under ABS and 20 under UP out of 45 groups; most proxies were not among the highest-ranked cells under either rule. The mean signed difference, log(rank*_UP_*) *−* log(rank*_ABS_*), was *−*0.104. The exact two-sided sign-flip analysis of the five nonzero paired differences gave *p* = 0.125; none of the five additional marker-rule contrasts against ABS survived Holm correction. These analyses do not establish a universally superior marker-selection rule or a unique causal cell type. In particular, the first-ranked heart-failure annotations under both ABS and UP did not pass within-family false-discovery correction, as was also the case for non-small-cell lung cancer. Figure 4 and Tables S2–S3 report the leader-level q-values and the six proxy ranks without treating the proxy as a causal ground truth.

**Figure 4:**
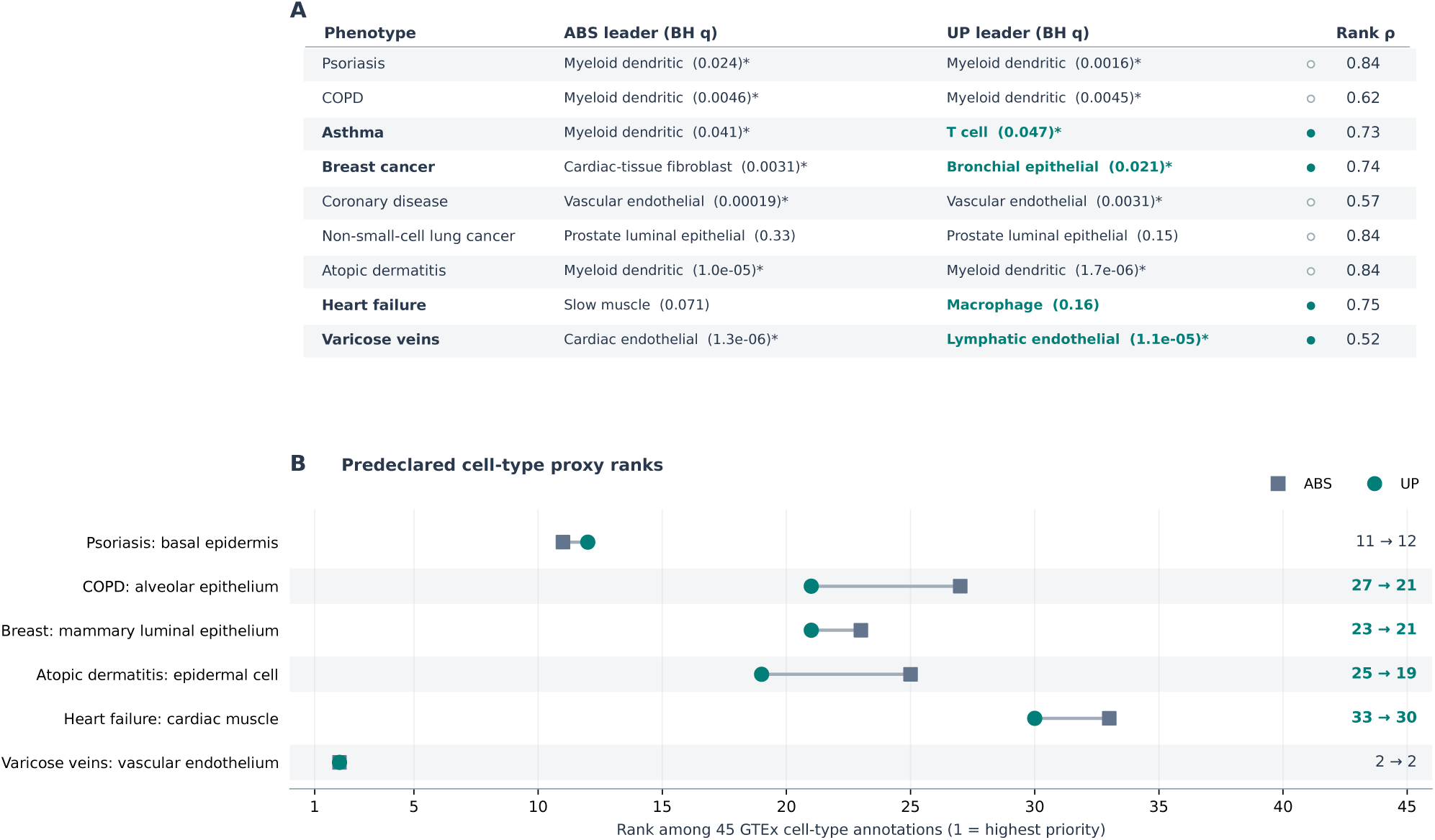
Marker-direction sensitivity in nine FinnGen release 13 genetic phenotypes. (A) Leading cell annotation under absolute-fold-change (ABS) and up-regulated (UP) marker rules, with corresponding within-rule Benjamini–Hochberg adjusted *q* values in parentheses. An asterisk marks *q <* 0.05; filled symbols denote different first-ranked annotations. Spearman *ρ* summarizes rank concordance over the 45 annotations shared between these families. The first annotation changed in four phenotypes, but only two shifts crossed broad cellular compartments (breast cancer and heart failure); neither heart-failure leader had *q <* 0.05. (B) Ranks of six broad cell-type proxies recorded before these cell-rank analyses. Lower ranks indicate higher priority. ABS is shown by squares and UP by circles. The proxy analysis is exploratory; the paired two-sided exact sign-flip test gave *p* = 0.125 for five nonzero differences. No claim of causal-cell identity or statistical independence across phenotypes is made.

### 3.6 The same nine stromal loci carry three different cell labels

We then checked whether the rank changes were driven by different genomic loci. We identified 57 independent genome-wide significant KOLLER_EUR loci. Of these, 52 had at least one eligible expressed candidate gene. Thirty-two were assigned to a specific compartment (stroma 9, endothelium 8, immune 6, epithelium 5, and perivascular/smooth muscle 4); another 20 were classified as broadly expressed, including regions containing FSHB, ESR1, CDKN2A/B, and GNRH1. Five loci lacked an eligible expressed candidate gene. Stromal loci were nominally enriched against the genomic background (*p* = 0.03), but no single locus determined the overall cell-state ranking. Removing genes near all 57 loci removed the bidirectional venous association (*p* changed from 3 *×* 10*^−^*^5^ to 0.14), whereas the up-only decidual association weakened but remained nominal (*p* changed from 1.4 *×* 10*^−^*^4^ to 0.021).

Removing the nine stromal loci (WNT4, GREB1, IGF1 region, TBX3, WT1, TSHZ3, HOXA9– 11, OSR2 region, SFRP1) was compared with removing 200 random 9-locus subsets (Figure S6). It weakened three signals more than chance: the up-only decidual set (+1.57 log10 units; permutation p = 0.005), the down-only luminal and ciliated sets (+1.69 and +1.35; p = 0.005), and the primary *|* log_2_ FC*|* venous set (+1.35 vs a random median of +0.69; p = 0.020). Removing the eight endothelial loci affected the venous set no more than chance (+0.89). One stromal genetic signal therefore appears as “decidual stroma” under up-only markers, “venous endothelium” under *|* log_2_ FC*|* and “luminal or ciliated epithelium” under down-only markers. Two secondary vascular *|* log_2_ FC*|* sets followed only partly (Arterial/CTL p = 0.070; Venous/ENDO p = 0.318). This test was formulated after the locus atlas showed the pattern and is labelled post hoc.

Classifying the loci a second time with the independent GSE266265 atlas [12] gave the same picture: 82% agreement (Cohen’s κ = 0.77), with 7 of the 9 stromal loci stromal again (PARPBP and VPS13B became broad). This removes most of the circularity of the test above, because the loci are now classified in one atlas and tested in the other. Loci called stromal in GSE266265 carried the HECA up-only decidual set (p 1.4×10*^−^*^4^ to 2.3×10*^−^*^3^; +1.22 log10 units vs a random median of +0.18; permutation p = 0.005). The nine HECA-stromal loci carried the GSE266265 eF3 set (p 8.0 × 10*^−^*^5^ to 7.6 × 10*^−^*^4^; +0.98 vs +0.13; permutation p = 0.005) (Figure S3).

The genes at these loci are expressed mainly in proliferative, hormone-responsive and decidual endometrial stroma (Figure S4). For example, SFRP1 reaches 2,023 CPM in eStromal; WT1 peaks in eStromal, eStromal_cycling and dStromal_early; WNT4 in eStromal_MMPs, sHormones and dStromal_mid. Endometriosis loci also overlap strongly with leiomyoma loci: 46% of loci vs 10% of matched random windows (OR 7.7, p = 4 × 10*^−^*^10^). Overlap with type 2 diabetes (OR 1.9) and hypertension (OR 2.3) was weaker.

### 3.7 Endometriosis risk preferentially maps to hormone-responsive endometrial stroma; evidence on mesothelium is inconsistent

To check whether the stromal result depended on the HECA atlas, we turned to GSE266265 [12], which is not part of HECA. This dataset includes 60 donors and 34 eligible cell types, analyzed with donor-paired pseudobulk. We compared four candidate cellular contexts often discussed in endometriosis: endometrial stroma, perivascular or mesenchymal stem-like cells, epithelium, and mesothelium. These comparisons assess where risk-associated genes are expressed; they do not by themselves determine the cellular origin of a lesion (Figure S7).

With size-matched up-only markers, endometrial stromal fibroblasts ranked first for all three endometriosis GWAS (overlapping samples, not independent replications): eF3 for KOLLER_EUR (p = 7 × 10*^−^*^5^, q < 0.01) and GCST90205183 (p = 1 × 10*^−^*^4^), and dS3 for FinnGen (p = 8 × 10*^−^*^3^). For KOLLER_EUR the best perivascular type ranked 2nd, the best epithelial type 11th and endothelium 13th. eF3 enrichment was larger for endometriosis than for type 2 diabetes (p = 0.013), hypertension (p = 0.029) and leiomyoma (p = 0.008), with the covariance between GWAS estimates taken into account. The hypertension control placed arterial endothelium and vascular smooth muscle first (p = 1 × 10*^−^*^5^). The thresholded rules did not replicate the HECA flip here: up-only ranked vascular smooth muscle first, and *|* log_2_ FC*|* gave only a weak immune signal (cDC1, q = 0.12).

#### Donor stability

We recomputed the donor-paired DE with each of the 60 donors left out. eF3 stayed enriched for KOLLER_EUR in every run (p ≤1.0 × 10*^−^*^3^; standardized coefficient 0.026–0.035). The stromal group stayed first in 48 of 60 runs for KOLLER_EUR, 60 of 60 for GCST90205183 and 54 of 60 for FinnGen. The prespecified threshold (90% for both main GWAS) was therefore missed for KOLLER_EUR: in the other 12 runs perivascular Prv-MYH11 ranked first. Those runs left out different donors, so for KOLLER_EUR stroma and perivascular cells are close rather than separated by one influential donor.

#### Identity of the stromal types

eF3 is a progesterone- and estrogen-receptor-positive fibroblast state with early decidualization (PGR 464, ESR1 243 and IGFBP1 405 CPM, plus PRLR) and few perivascular markers (SUSD2 4, MCAM 31 CPM). dS3 is decidualized stroma (PTGDS, APOD, CRISPLD2; IGFBP1 1,222 CPM). Genes at the stromal loci are stromal rather than perivascular: WT1 160–200 vs 26–39 CPM, and IGF1 1,256–2,449 vs 283–722 CPM.

#### Stroma versus perivascular cells

In joint models, eF3 kept p = 1–3×10*^−^*^4^ given each of five perivascular or mesothelial-labelled sets, and dS3 kept p < 0.05 in all three GWAS. Perivascular sets also survived given eF3 for KOLLER_EUR, so these are two separable programs; the stromal one is stronger and trait-specific. eF3 survived conditioning on mean expression, breadth and LOEUF (p 6.7 × 10*^−^*^5^ to 2.5 × 10*^−^*^3^).

#### Mesothelium

The cell type labeled “mesothelial” in GSE266265 did not have the expected marker profile: its MSLN expression was 14 CPM (compared with 27 in glandular epithelium), and UPK3B was only 2 CPM. We therefore did not treat that label as reliable evidence about peritoneal mesothelium. Instead, we identified a more stringent mesothelial population from the raw GSE213216 dataset [13]: 1,069 cells across 12 samples, with MSLN 549, UPK3B 416, and ITLN1 2,048 CPM, but no EPCAM or PTPRC. For KOLLER_EUR, mesothelium ranked third, behind lesion stroma (first, *p* = 0.05) and eutopic stroma (second); unaffected peritoneal fibroblasts ranked sixth. The endometriosis associations were nominal, whereas the hypertension control strongly favored perivascular and smooth-muscle cells (*p* = 1 *×* 10*^−^*^5^). This result did not establish mesothelial enrichment, but it did not exclude a mesothelial role.

A second lesion cohort gave a different answer. Applying the same selection rule to GSE179640 [14] identified 779 mesothelial cells in 14 samples from peritoneal lesions and nearby peritoneum. Under size-matched up-only markers, mesothelium ranked first for all three endometriosis GWAS (KOLLER_EUR *p* = 0.018; GCST90205183 *p* = 0.09; FinnGen *p* = 0.0046, *q* = 0.04), but last or nearly last for hypertension, leiomyoma, and type 2 diabetes. Continuous specificity instead favored peritoneal-lesion stroma (all three endometriosis *p* = 0.02–0.07). Thus the prespecified expectation that endometrial-type stroma would outrank mesothelium was not met in this cohort. In follow-up analyses, mesothelial associations persisted after conditioning on stromal sets (KOLLER_EUR *p* = 0.022; FinnGen *p* = 0.006), and after removing all 52 eligible KOLLER_EUR locus genes (*p* = 0.051 and 0.016); four of those genes, including WT1, were among the mesothelial markers. Because the cohorts were small, their sampling differed, and the selection rules disagreed, we regard the evidence on mesothelium as unresolved.

#### Inside lesions

We asked a separate question: do genes near the stromal GWAS loci remain enriched in stromal cells when those cells are found inside endometriosis lesions? Using donor-averaged lineage pseudobulk in GSE213216, we compared each candidate gene’s expression in its assigned lineage with expression in the other cells of the same sample. We then compared the mean difference with 10,000 expression-matched random gene sets. The nine stromal-locus genes were enriched in stroma from peritoneal lesions (12 donors; mean log_2_ ratio 2.48, versus a null 95th percentile of 0.86; permutation *p* = 1 *×* 10*^−^*^4^), from endometriomas (6 donors; 1.38 versus 0.44; *p* = 2 *×* 10*^−^*^4^), and from eutopic endometrium (9 donors; 2.24 versus 0.46; *p* = 1 *×* 10*^−^*^4^; Supplementary Figure S8). All nine genes were individually more highly expressed in stroma than in the rest of peritoneal-lesion samples (one-sided *p <* 0.05), and six of nine were in endometriomas. Stroma was the highest-expressing lineage for 7/9 genes in eutopic endometrium, 5/9 in peritoneal lesions, and 4/9 in endometriomas. WNT4, GREB1, and TSHZ3 peaked in stroma in all three tissues, whereas WT1 and PARPBP were highest in mesothelium and TBX3 and HOXA9 in perivascular cells in peritoneal lesions.

The expected lineage specificity of endothelial and immune locus genes was retained in both lesion types (permutation *p* = 1 *×* 10*^−^*^4^). Epithelial-locus genes passed the test in peritoneal lesions (*p* = 3 *×* 10*^−^*^4^) but not in endometriomas (*p* = 0.07); the three perivascular-locus genes did not pass in either. Only two donors contributed unaffected peritoneum in this cohort, preventing the intended comparison with host peritoneal fibroblasts.

GSE179640 offered a second look, with eight donors contributing peritoneal lesions and four endometriomas. Stromal-locus genes were enriched in the stromal lineage in both tissues (mean log_2_ ratios 2.66 and 1.60; permutation *p* = 1 *×* 10*^−^*^4^ for each). Importantly, the same genes were also enriched in stroma from lesion-adjacent peritoneum (2.53). Within six paired donors, lesion stroma did not exceed adjacent-peritoneum stroma (difference 0.01 log_2_ units; *p* = 0.26). The prespecified H-HOST prediction therefore failed. These findings localize the genes to fibroblast-like cells, including host peritoneal fibroblasts; they do not reveal whether those cells originated in the endometrium.

#### Non-European GWAS

We also examined summary statistics released by Koller et al. for East Asian, African, and admixed American participants. The effective sample sizes were approximately 8,700, 9,500, and 6,700, respectively, compared with approximately 198,000 for the European analysis. Using ancestry-matched 1000 Genomes reference panels, the eF3 stromal program did not show nominal enrichment under size-matched up-only markers (East Asian: *p* = 0.076, rank 6/34; African: *p* = 0.74; admixed American: *p* = 1.0). The prespecified East Asian eF3 replication criterion was therefore not met.

Another endometrial fibroblast state, eF1, ranked first in the East Asian GWAS (*p* = 0.01, *q* = 0.40), whereas mid-secretory epithelium and lymphatic endothelium led in the African and admixed American analyses. These smaller GWAS had only 3–5% of the European effective sample size and produced 2, 0, and 0 genome-wide significant MAGMA genes, respectively, compared with 72 in the European analysis. Low power may explain the lack of replication, but it would be misleading to claim that the eF3 finding has been demonstrated across ancestries.

#### Other routes

An RNA-based polygenic scoring approach also supports stromal states. scDRS [15] on GSE213216 placed the strongest endometriosis signal in endometrial-type stromal states, in lesions (eF2, eF3, dS4) and eutopic tissue (dS2, eF3), q = 0.02. Fallopian-tube epithelium [16] showed no preference (tubal and uterine stroma were indistinguishable). Lesion macrophages, reported by Ochoa et al. [7], were not reproduced under our reconstruction; their donor-level excess was shared with hypertension and type 2 diabetes.

### 3.8 Most HECA states split by disease status come from one or two donors, so the stromal compartment is robust but a single stromal state is not

A cell-state label can appear robust even when almost all its cells come from one donor. That is a particular concern when cell types are split by disease status. In HECA, the largest donor contributed more than 80% of the cells in 16 of 45 status-specific groups and more than half in 32 of 45. The median effective number of donors per group was only 2.3 (2.0 in the count-matched cohort). The decidual groups were particularly concentrated: one donor supplied 97% of dStromal_mid/ENDO, another supplied 92% of dStromal_early/ENDO, and a third supplied 89% of dStromal_early/CTL. A ranking of those groups can therefore partly reflect who happened to contribute cells (Figure 5a).

**Figure 5:**
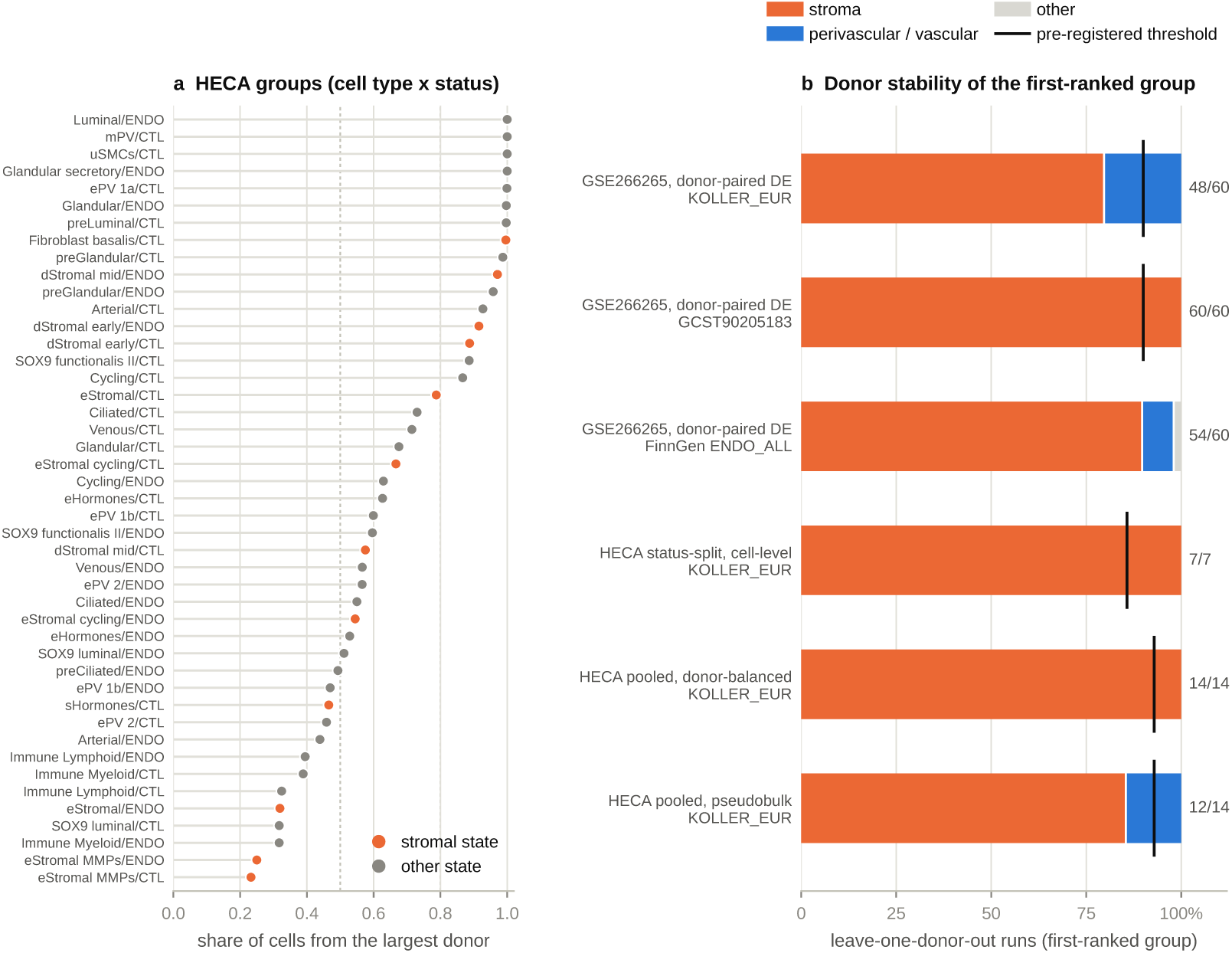
Donor structure and donor stability. (a) Share of cells contributed by the largest donor in each HECA cell type × disease-status group with ≥30 cells (the design of Koller et al.); stromal states in orange. (b) Leave-one-donor-out stability of the first-ranked group (MAGMA, size-matched up-only sets): GSE266265 donor-paired DE (60 donors; three endometriosis GWAS) and three HECA designs for KOLLER_EUR (status-split labels with cell-level DE, 7 endometriosis donors left out; labels pooled over status with ≤50 cells per donor and type, 14 donors; donor-paired pseudobulk with pooled labels, 14 donors). Right, runs with a stromal state first; black ticks, prespecified thresholds (for the status-split design the rule concerned decidual states, met in 6 of 7).

We tested donor dependence in three designs (MAGMA, KOLLER_EUR, size-matched up-only sets; Figure 5b):

- Leave one endometriosis donor out (status-split labels, cell-level DE). A decidual state ranked first in 6 of 7 runs and a stromal state in 7 of 7. Without the donor behind dStromal_mid/ENDO that state dropped out and dStromal_early/ENDO ranked first.
- Labels pooled over status, at most 50 cells per donor and type (14 donors). A stromal state ranked first in the full data (eStromal_MMPs p = 2×10*^−^*^5^, eStromal 1×10*^−^*^4^, dStromal_mid 2 × 10*^−^*^4^) and in 14 of 14 leave-one-donor-out runs; perivascular states never ranked above stroma. With thresholded up-only sets dStromal_mid ranked first in 14 of 14. Controls behaved as expected (hypertension: arterial and venous endothelium; leiomyoma, thresholded up-only: perivascular ePV_1b). For GCST90205183 this design ranked perivascular ePV_2 first (p = 8 × 10*^−^*^3^).
- Donor-paired pseudobulk DE with pooled labels (up to 14 donors). dStromal_early ranked first (p = 9 × 10*^−^*^4^), and a stromal state stayed first in 12 of 14 runs, one short of the prespecified 13. The two exceptions left out the two donors who carry the decidual states; perivascular ePV_1b then ranked first. With status-split labels (3–7 donors per state) donor-level DE had little power (best p = 0.02, perivascular).

Across designs the stromal compartment is robust. Which stromal state ranks first (decidual, proliferative or early-decidualizing) depends on the marker rule and on which donors are included.

### 3.9 Chromatin accessibility supports a stromal regulatory context in the initial tests

The RNA analyses suggested a broad stromal program, but they all began with gene expression. To test that interpretation in a different molecular layer, we examined the accessible chromatin of GSE183771 [17]. Its 13 single-nucleus ATAC libraries correspond to 12 endometrial biopsies collected on cycle days 7–25. The specimens came from women attending a reproductive clinic and were not selected as an endometriosis case–control series. We retained 36,073 nuclei (36,177 were reported in the source study), merged library-specific peak calls into 293,258 regions, and mapped the accessible regions to GRCh37. Of the autosomal peaks, 99.8% were lifted successfully. We selected accessible peaks by biopsy-paired lineage specificity, *not* by RNA differential-expression gene lists. A fixed panel of 46 marker genes was nevertheless used to assign the ATAC cell-cluster labels; the entire analysis is therefore not marker-free.

Before opening GWAS data, the chromatin pipeline was required to pass four checks covering genome mapping, lineage composition, representation across biopsies, and overlap with cultured endometrial stromal-cell ATAC. The first two clustering-label designs failed the composition check and were revised before any GWAS was read. The final labels were close to the published compartment proportions (mesenchymal 46.7% versus 46.6%; epithelial 42.7% versus 45.6%; immune 6.1%; endothelium 1.3%; unassigned 3.2%). Some mesenchymal nuclei could not be reliably distinguished as stroma or perivascular cells; no confident perivascular cluster was identified. After ranking 240,290 eligible peaks, we retained the top 10% for each lineage. An independent annotation check was reassuring: 67% of stromal-specific peaks overlapped accessible regions in cultured stromal cells (GSE246585), compared with 32% of epithelial-specific and 24% of all peaks (Figure 6b).

**Figure 6:**
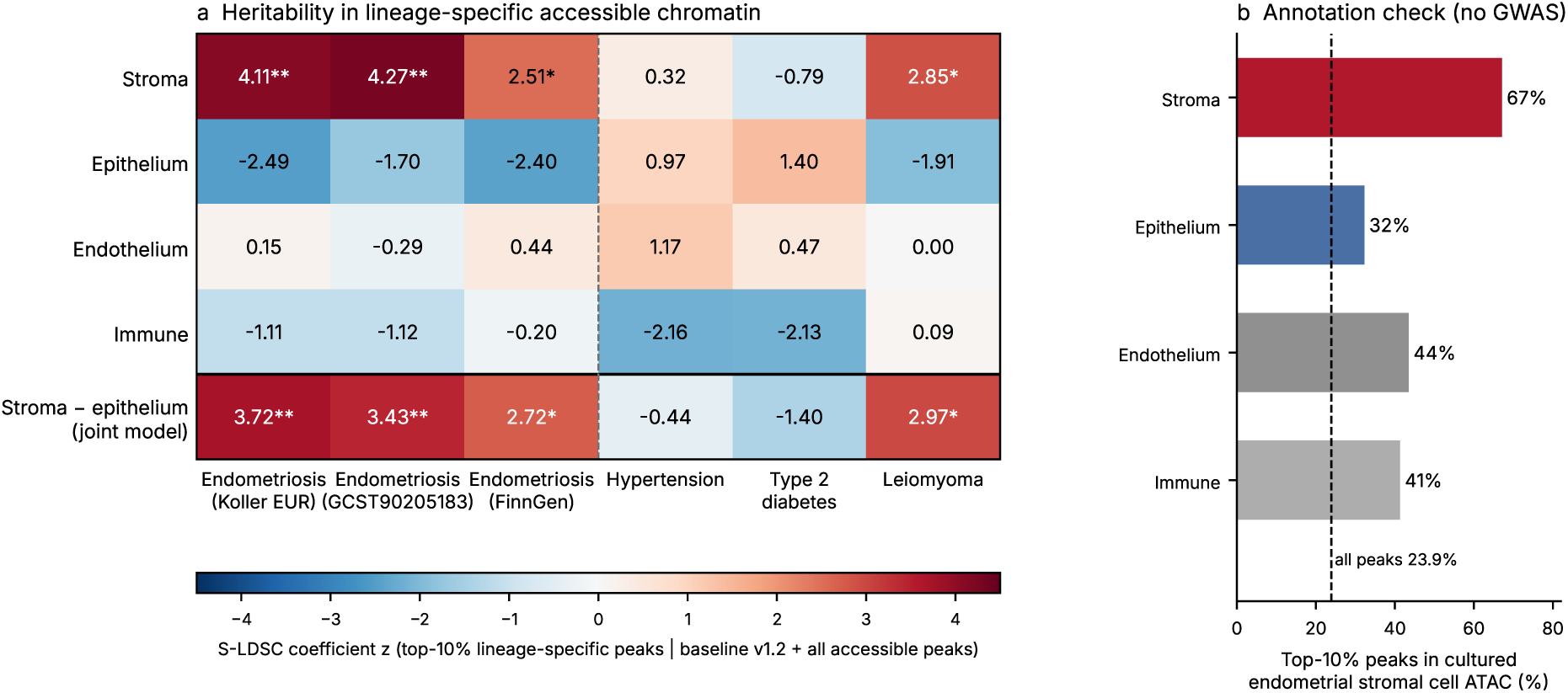
Chromatin-accessibility enrichment independent of RNA-DE marker-set selection (GSE183771). (a) S-LDSC coefficient z of lineage-specific accessible chromatin (top 10% of peaks by biopsy-paired specificity), with all accessible peaks and baseline v1.2 in the model; bottom row, stroma minus epithelium in one model. * one-sided p < 0.05; ** p < 0.001. The three endometriosis GWAS overlap in samples; hypertension and type 2 diabetes are control traits; leiomyoma is descriptive. (b) Share of each lineage’s top-10% peaks that overlap peaks of cultured endometrial stromal cells (GSE246585); dashed line, all peaks. This check was fixed and run before any GWAS file was read.

With the annotations fixed, we ran the three initially specified chromatin tests once (S-LDSC with baseline v1.2 and all accessible peaks as control; Figure 6a). Stroma-specific chromatin was enriched for KOLLER_EUR (z = 4.11, one-sided p = 2 × 10*^−^*^5^; CH-1). It was more enriched than epithelium-specific chromatin in one model (z = 3.72, p = 1 × 10*^−^*^4^; CH-2), and stroma ranked first among the four lineages (stroma 4.11, endothelium 0.15, immune −1.11, epithelium −2.49; CH-3). The result held for the two other endometriosis GWAS, which overlap KOLLER_EUR in samples (GCST90205183 z = 4.27; FinnGen z = 2.51, p = 0.006), and for top-5% and top-20% peak sets (z = 3.40 and 4.01). Epithelium-specific chromatin was depleted relative to all accessible peaks in all three endometriosis GWAS. Hypertension (z = 0.32) and type 2 diabetes (z = −0.79) showed no stromal enrichment, so the result is negative for the two unrelated traits under the originally registered rule. Leiomyoma, a smooth-muscle tumour of the uterus, also ranked stroma first (z = 2.85, p = 0.002), which suggests a regulatory program shared across uterine mesenchyme; no smooth-muscle annotation was available to separate the two. In the Koller cohort the *|* log_2_ FC*|* rule ranked venous endothelium first; endothelium-specific chromatin showed no enrichment (z = 0.15), although this annotation was weakly specific and rested on 458 nuclei.

### 3.10 Run-specific hypotheses, sequential follow-ups, and audit outcomes

Tests fall into five groups, all reported (Table S1). Original confirmatory tests were fixed before any related result was seen: 7 tests, 4 supported and 3 not. Follow-up tests were formulated after an earlier result and registered before their own run: 25 tests, 13 supported (one only nominally), 11 not, and 1 that met its rank criterion with nominal signals only. Audit tests answered an internal review of the draft and were registered before their runs: 7 tests, 4 supported and 3 not. The chromatin tests were registered after a GWAS-free audit of the annotations and before any GWAS file was read: 3 tests, all supported. Descriptive steps are listed for context. Many tests share GWAS and atlases, so the counts are not independent confirmations. The expanded nine-phenotype analysis was tracked separately: a descriptive change-of-leader endpoint, an exploratory six-proxy paired test, and five exploratory rule-vs-ABS comparisons with Holm adjustment. These later analyses are included in the supplementary analysis registry and are not added retrospectively to the earlier original-test counts. Three earlier audits (decidual specificity by LDSC-SEG, the lesion-macrophage result and the fallopian tube) are summarized in the separate Supplementary Note.

### 3.11 Joint chromatin models and sensitivity to individual biopsies

The original scATAC tests analyzed each lineage annotation separately. After seeing those results, we asked a harder question: does the stromal signal persist when competing lineages are considered together? We tested it in a model that included the all-accessible-peaks control and stroma, epithelial, endothelial, and immune annotations *simultaneously*. Because these models were planned after the initial association was observed, they are reported as sensitivity analyses, not as independent prespecified confirmations. Chromatin peaks were selected by differences in accessibility across biopsies, rather than by RNA differential-expression sets. RNA marker genes were used to label cell clusters, however, so this is not a completely marker-free analysis. In KOLLER_EUR, the stromal coefficient remained positive after joint adjustment with baseline v1.2 (*z* = 3.891, one-sided *p* = 4.995 *×* 10*^−^*^5^) and baselineLD v2.2 (*z* = 3.828, *p* = 6.471 *×* 10*^−^*^5^). The covariance-aware contrast between stromal and epithelial coefficients was also positive in both models (*z* = 3.781, *p* = 7.81 *×* 10*^−^*^5^ and *z* = 3.677, *p* = 1.182 *×* 10*^−^*^4^, respectively). As summarized in Figure 7 and Table S4, with baselineLD v2.2, stromal coefficients were positive for two additional, partly overlapping endometriosis GWAS (GCST90205183: *z* = 4.055; FinnGen ENDO_ALL: *z* = 1.979), and for leiomyoma (*z* = 2.418), but not for hypertension (*z* = 0.177) or type 2 diabetes (*z* = *−*0.797). Thus the signal was not restricted to endometriosis; a uterine mesenchymal regulatory context may be shared with leiomyoma. The additional GWAS do not constitute independent genetic replication.

**Figure 7:**
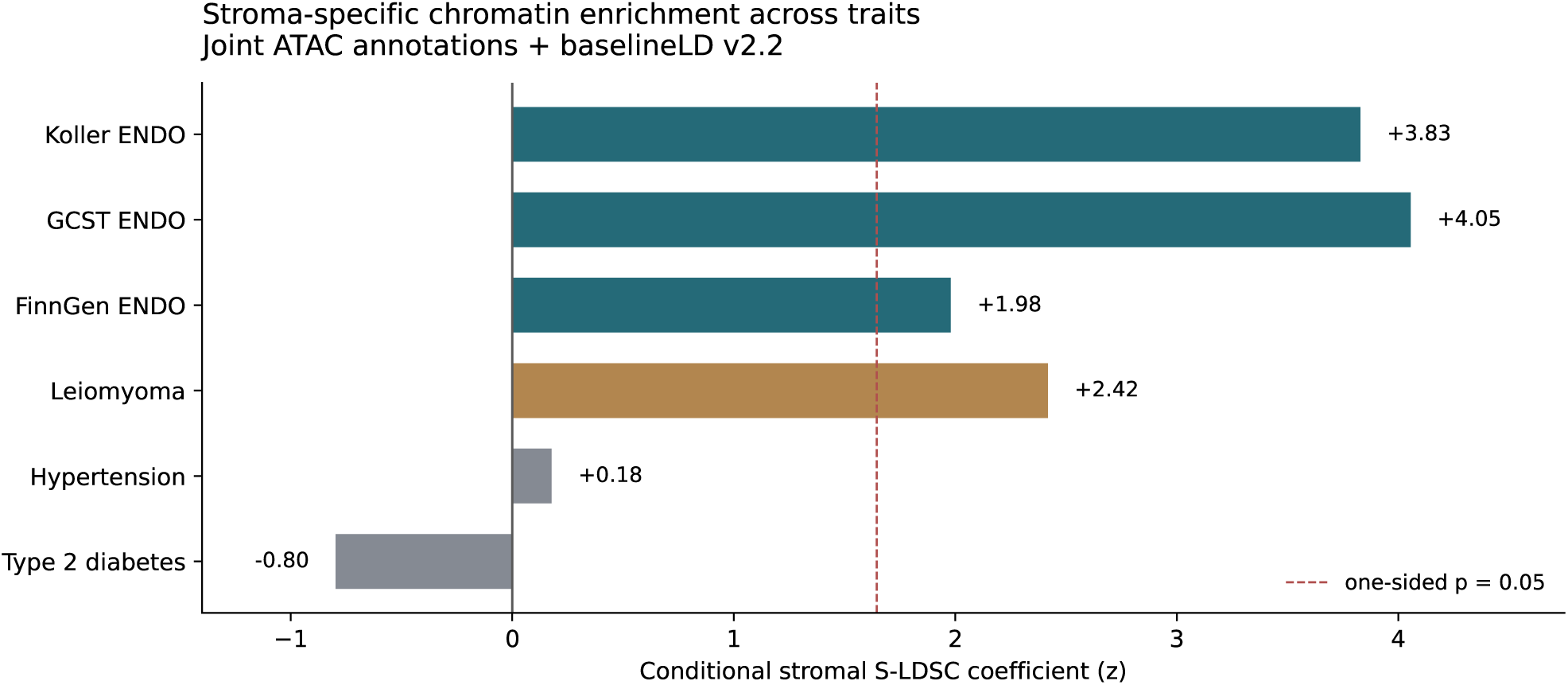
Post-result full joint chromatin models. Conditional S-LDSC *z* statistics for the stromal accessibility annotation in a joint model that includes all accessible peaks and four lineage-specific top-10% annotations, alongside baselineLD v2.2. The three endometriosis GWAS overlap in participants and are not independent genetic replications. Leiomyoma is a related uterine comparator; hypertension and type 2 diabetes are negative-comparator traits. One-sided nominal tests; this analysis followed, rather than preceded, the initial CH-1–CH-3 findings.

We then asked whether one influential biopsy could explain the joint result. The two technical libraries from biopsy S19a_18189 were pooled as one unit. Using the original cell labels and fixed accessible-peak universe, we reconstructed the full-data peak rankings exactly: all four top-10% selections and all five 1000 Genomes SNP annotation masks matched the frozen original annotations, and the complete joint regression numerically reproduced the preceding full-data joint fit before exclusions began. We then recalculated peak specificity, SNP annotations, LD scores, and the KOLLER_EUR joint regression after excluding one biopsy at a time. Every one of the 12 refits retained positive nominally significant stromal enrichment (conditional *z* = 3.439–3.890, one-sided *p* = 5.01 *×* 10*^−^*^5^–2.92 *×* 10*^−^*^4^). The stromal-minus-epithelial contrast also remained positive in all 12 fits (*z* = 3.544–3.891, *p* = 4.99 *×* 10*^−^*^5^–1.973 *×* 10*^−^*^4^). The association remained detectable regardless of which one biopsy was removed. They do not supply 12 independent replications or account for uncertainty in the frozen cell clustering. Full results are given in Supplementary Tables S5–S6 and Supplementary Figure S9.

## 4 Discussion

Our central finding is straightforward: changing the definition of a marker gene can change the cell type that appears most relevant to a GWAS. In the count-matched Koller analysis, selecting genes by the magnitude of their fold change favored venous endothelium, whereas selecting only up-regulated genes favored decidual stroma. The latter RNA association was nominal (*q* = 0.08 after within-rule adjustment), so the rank difference should not be mistaken for two equally supported discoveries. The rules answer different questions. A bidirectional set includes genes a cell expresses relatively weakly; an up-only set focuses on genes more active in that cell. Our benchmark explains when that distinction matters, while the donor analyses show why naming one precise stromal state can be premature. Chromatin accessibility from a separate biopsy series provides convergent support for a broader stromal regulatory context. It does not prove that venous endothelium is irrelevant, especially given the sparse endothelial ATAC annotation.

Recommendations for marker-based enrichment.

1. Report the direction of the marker set and justify it. If the hypothesis is that risk genes are active in a cell type, use up-type or specificity sets.
2. Do not rely solely on thresholded *|* log_2_ FC*|* sets. In our benchmark the rule was never best, and in selected settings it missed a positive control found by an up-type rule. In a six-trait known-source panel it matched thresholded up-only markers (3 of 6 each), but no rule was best everywhere, so the choice should be checked against positive controls.
3. Include positive-control traits with known cell types (here leiomyoma, hypertension and four further traits), and a trait-contrast test against unrelated traits that accounts for overlap between GWAS.
4. Use joint models (a set given its competitors) before naming one cell type, and report related-state ambiguity: decidual sources were placed in the right stromal group far more often than in the exact state. Fine-mapping of cell-type annotations into credible sets formalizes this step [10].
5. Check the top hit at the locus level: which loci carry it, and do they belong to the named cell type in an independent atlas?
6. Check donor composition before naming a cell state: report the effective number of donors per state, and repeat the ranking with donor-balanced or donor-level DE and with donors left out.
7. Where possible, test the conclusion in another molecular layer without reusing the same RNA marker-selection rule, such as cell-type-specific accessible chromatin from other donors.

The benchmark also sets an important limit on the claim. Mixing up- and down-regulated genes can *mask* a signal carried by genes that are active in the cell type; we observed this for decidual stroma and the leiomyoma smooth-muscle control, across S-LDSC and MAGMA. A second possibility is that a cell type looks falsely associated because of its down-regulated genes. That explained individual venous hits, but it was not a general property of all bidirectional sets. A related example of the gap between computational stability and biological interpretation was seen in a separate multi-omics benchmarking study [27]. Atlas composition matters: about 60% of bidirectional HECA markers were down-regulated, compared with a median of 33% in GTEx. In the known-source panel, their depleted share also did not separate successful assignments (6–37%) from failures (28–34%). We therefore recommend checking marker direction and relevant controls rather than claiming that one selection rule always wins.

### Biological interpretation

The results point to a consistent *compartment*, not one uniquely defined cell state. Under direction-aware gene selection, endometriosis risk was associated with hormone-responsive endometrial stromal programs in HECA, with eF3 and dS3 fibroblast states in GSE266265, and with lesion and eutopic stromal states in scDRS. An independent scATAC cohort provided convergent evidence from accessible chromatin. A recent pan-organ atlas similarly associated endometriosis genetic risk with endometrial fibroblast programs later in the menstrual cycle [19]. These sources are distinct at the cell-measurement level, but several analyses reuse overlapping GWAS participants and should not be counted as independent genetic replications.

The precise state was less stable than the broad lineage. Many disease-status-specific HECA groups were driven by one or two donors, and GSE266265 did not always distinguish eF3 from a perivascular alternative for KOLLER_EUR. Nine loci containing developmental or hormone-responsive candidates (WNT4, GREB1, WT1, HOXA9–11, IGF1, SFRP1, TBX3, TSHZ3, and the OSR2 region) underpinned much of the stromal association. Their broad stromal assignment was similar across two RNA atlases, but gene expression at a locus is not proof of which variant or cell population mediates disease risk.

A further caution comes from leiomyoma. It shares part of the stromal chromatin association with endometriosis, as well as locus-level overlap (OR 7.7 in the analysis described above). This points to a possible shared uterine mesenchymal program rather than an endometriosis-specific cellular mechanism. Other strong endometriosis loci, including FSHB, GNRH1, and ESR1, act in broader hormonal pathways. The stromal result is therefore one component of inherited risk, not a complete explanation of the disease.

### What this means for theories of origin

It is tempting to interpret stromal enrichment as evidence that endometriosis starts in one particular stromal cell. Our data cannot make that leap. They are compatible with models in which menstrual endometrium carries inherited risk, but do not distinguish the cells that initiate a lesion from the stromal cells that may later colonize it or be recruited from surrounding tissue. The two relatively small mesothelial datasets point in different directions. Mesothelium and endometrial stroma also share developmental features and markers such as WT1, making their separation difficult.

Somatic-mutation studies have found oligoclonal epithelial lesions with ancestry in normal endometrium, alongside genetically distinct stromal cells [20]. Cancer-associated mutations, including KRAS, PIK3CA, and ARID1A, can occur in lesion epithelium [21, 22]. These observations raise a possible division of roles between inherited risk and somatic changes, but do not establish one. In our own data, stromal-locus genes remained expressed in lesion-associated fibroblasts, yet were also enriched in nearby unaffected peritoneal fibroblasts. A direct test of origin would require matched genotype, somatic-mutation, single-cell or spatial information from the same patients. Our finding concerns the regulatory context in which inherited risk may act, not the first cell from which a lesion arises.

### What the chromatin sensitivity analyses add

The original change in RNA-based rankings was more than a matter of which annotation happened to have the smallest *p* value. A direction-aware interpretation repeatedly implicated endometrial stromal programs, but the identity of the highest-ranked stromal state depended on donors and on the atlas. The scATAC analysis provides a different view: in another set of biopsies, a stromal accessibility signature retained a positive coefficient after modeling all four available lineage annotations together and after conditioning on an expanded baseline of genomic features. The association also survived exclusion of every individual biopsy. This convergence strengthens the case for a broad stromal regulatory context, while leaving open which molecular state and which variants mediate the effect. It is important that RNA gene markers were still used to label the ATAC cells; this was a test independent of RNA marker *set selection*, not of all marker-based information.

### Boundaries of biological interpretation

The ATAC donor series was not recruited as an endometriosis case-control cohort, and ten of twelve biopsies were collected during the secretory phase. The endothelial cluster contained only 458 nuclei and no reliable perivascular cluster was resolved. We therefore cannot distinguish endometrial stromal fibroblasts from every related mesenchymal state, nor infer a causal cell of lesion origin. Stromal chromatin also carried a leiomyoma signal, consistent with some shared uterine mesenchymal regulation. The three endometriosis GWAS overlap in participants and do not represent three independent genetic validations. Likewise, the 12 LOO estimates are highly correlated and must not be treated as 12 independent confirmations. These limits do not negate the observed conditional enrichment, but they narrow the claim to a regulatory context for a portion of inherited risk.

### Cross-trait interpretation

The changed leading annotations across several phenotypes demonstrate method sensitivity rather than biological ground-truth accuracy. Some leading annotations are drawn from anatomically distant tissues, and several rank-one associations do not survive within-family false-discovery correction. The six reference proxies were broad biological hypotheses, not independently proven causal cell types. The small set of correlated GWAS, shared participants, atlas composition, and the inability to exclude unarchived pre-amendment exploratory outputs limit causal and confirmatory interpretation.

### 4.1 Limitations

- No independent genetic replication. KOLLER_EUR overlaps heavily with GCST90205183 and FinnGen ENDO_ALL (rg 0.96 with each; LDSC cross-trait intercepts 0.53 and 0.69). GCST90205183 and FinnGen R13 are closer to independent (rg 0.89, intercept 0.05); GCST90205183 includes an earlier FinnGen release. FinnGen is the weaker GWAS. Trait contrasts account for the covariance between GWAS estimates through paired genome rotations. The non-European GWAS of Koller et al. are independent of these samples but much smaller, and they did not replicate the eF3 signal.
- Modest effect sizes. The S-LDSC up-only decidual signal in the exact cohort is nominal (q = 0.08). Trait contrasts reach one-sided p of 0.01–0.04 and are not corrected across the two atlases or the secondary contrasts. Positive controls in S-LDSC are nominal only; MAGMA carries the control evidence.
- Donor numbers. HECA states split by disease status rest mostly on one or two donors, and donor-level DE in HECA had little power. Two prespecified donor-stability thresholds were missed narrowly (GSE266265 KOLLER_EUR 48 of 60 runs; pooled HECA pseudobulk 12 of 14). We therefore claim the stromal compartment, not a specific stromal state.
- Thresholded rule flip not general. It did not replicate in GSE266265, where the thresholded up-only rule ranked vascular smooth muscle first. It was also threshold-sensitive in HECA (decidual first in 5 of 8 settings; stromal in 8 of 8).
- Post hoc steps. Several hypotheses were formulated after earlier results. They were registered before their own runs, and several were not supported. One fixed rule was mis-specified (the GTEx “contractile cell” label check); we report both the rule’s output and the corrected label. The stromal-locus test is conditional on an expression-based classification of loci; the cross-atlas version reduces this circularity but does not validate a mechanism.
- Mesothelium. Real mesothelial cells were few in both lesion cohorts (1,069 cells in 12 samples; 779 in 14), the two cohorts disagreed, all signals were nominal except one (q = 0.04), every GSE179640 patient was hormonally treated, and samples from one patient are not independent.
- Known-source panel. Six traits in one atlas; the expected cell types rest on prior knowledge and on label rules, two of which were corrected after the label list was seen. Thresholded *|* log_2_ FC*|* and up-only markers tied (3 of 6), so the panel does not show that up-only markers are generally better.
- Lesion samples. GSE213216 provides 12 donors with peritoneal lesions, 6 with endometriomas and 2 with unaffected peritoneum; GSE179640 adds 8, 4 and 6 (lesion-adjacent peritoneum). Lesion stroma is defined by markers and mixes endometrial-type and host fibroblasts, and expression in a lineage does not show which cells carry the risk alleles.
- Dataset overlap and quality. GSE213216 is part of HECA; GSE266265 is not. GSE266265 counts are non-integer (ambient-corrected); CPM-based analyses are unaffected. DE methods differ by dataset (cell-level Wilcoxon in HECA and GTEx, donor-paired pseudobulk in GSE266265 and GSE213216); HECA was also analysed with donor-balanced and donor-paired designs. A separate audit of endometriosis microarray classifiers identified exact reuse of CEL files across nominally distinct GEO series and outcome-informed batch correction [28]. That study addressed diagnostic prediction, not GWAS-to-cell-state enrichment, and provides no independent validation of the present findings.
- Chromatin test. GSE183771 has 12 donors, mostly in the secretory phase (10 of 12), from a reproductive clinic (0–7 previous miscarriages) and not selected for endometriosis. Peaks were called per library and merged, so rare lineages are under-represented and the endothelial and immune annotations were weakly specific. Stroma could not be separated from perivascular cells, and leiomyoma shared the stromal signal; the result places heritability in stromal (uterine mesenchymal) regulatory regions, not in a specific stromal state. The cell labels were revised twice for technical failures before any GWAS was read.
- Interpretation. Enrichment shows where risk genes are active, not causal cell types or where lesions start. Related endometrial states are hard to separate; we report group-level and exact-state recovery separately.

## Author contributions

E.K. conceived the study, designed and performed the computational analyses, organized the data and reproducibility record, and drafted the manuscript. A.A. provided critical scientific feedback and reviewed and edited the manuscript.

## Acknowledgements

We thank the participants and investigators of the FinnGen study, and the authors of HECA, GTEx v9 snRNA-seq, GSE266265, GSE213216, GSE179640, GSE183771 and GSE246585 for making their data public. We thank Rahmioglu et al. and Koller et al. for releasing GWAS summary statistics.

## Use of AI tools

Large language models (Claude, Anthropic; ChatGPT, OpenAI) were used to draft text and to review the manuscript. The computational analyses were executed and audited by E.K. The authors are responsible for evaluating the accuracy of the analyses and scientific reporting; artificial-intelligence systems are not authors.

## Data availability

All inputs are public: GWAS summary statistics (GCST90205183 from the GWAS Catalog, the European, East Asian, African and admixed American summary statistics of Koller et al. from Zenodo, doi:10.5281/zenodo.18983492, and FinnGen R13, available via the official FinnGen release portal (https://www.finngen.fi/en/access_results)); atlases (HECA via E-MTAB-14039 and the HECA portal, GTEx v9 snRNA-seq via CELLxGENE, GSE266265, GSE213216 and GSE179640 via GEO; single-cell and cultured-cell chromatin GSE183771 and GSE246585 via GEO; 1000 Genomes superpopulation panels via Zenodo, doi:10.5281/zenodo.6614170); and resources (MAGMA reference files, LDSC baseline v1.2 and 1000 Genomes Phase 3 files, gnomAD v2.1.1 constraint). For the nine-phenotype extension, the fixed phenotype roster, dated proxy-label addendum, unchanged marker-definition code, full cell-type ranks, per-trait adjusted values, and analysis summaries are supplied as accompanying Supplementary Code and Data. The source archives carry checksums and execution logs to document the sequence of analyses. The author-maintained project repository is https://github.com/Helen2813/paper9. Scripts for the chromatin reconstruction, joint regression, and leave-one-biopsy-out sensitivity calculations accompany the versioned analysis materials. The accompanying code archive contains these scripts, the cross-trait analysis scripts, frozen definitions, and the result tables necessary to audit the added cross-trait claims. The large reference datasets and third-party raw matrices are not redistributed. Third-party datasets should be obtained from the original providers in accordance with their access and reuse conditions.

