## supplement file for "Marker direction alters genetic cell-state prioritization across complex traits: an endometriosis and chromatin case study"

### Supplementary note: earlier audits

**Decidual specificity by LDSC-SEG (GSE266265).**

Before the marker-direction analyses we tested whether decidual stromal types of GSE266265 are enriched for GCST90205183 with LDSC-SEG (control-donor marker sets, one-sided, Holm over four decidual types). dS4 ($p = 0.015$, Holm 0.061) and dS3 ($p = 0.029$, Holm 0.085) ranked 1st and 2nd of 42 types, but neither passed the prespecified Holm threshold, so the check was not replicated. No type passed BH in any trait (best $q = 0.60$), and the hypertension control placed perivascular and vascular smooth-muscle types at ranks 1–4 with $q = 0.26$: the within-tissue LDSC-SEG design was power-limited.

**Lesion macrophages (Ochoa et al.).**

Ochoa et al. [7] reported that M2 macrophages in endometriosis lesions carry germline risk (scDRS on GSE213216). We re-ran scDRS on 74,310 lesion cells, following their sample and immune re-annotation as closely as public metadata allowed. Macrophage groups did not pass Holm-corrected Monte Carlo association for either endometriosis GWAS (GCST90205183 Holm 0.365; KOLLER_EUR 0.739), so the claim was not reproduced under our reconstruction. A donor-consistent excess of disease-gene expression in lesion macrophages was present for endometriosis (myeloid $p = 3 \times 10^{-7}$, 29 of 30 donors) but equally for hypertension ($p = 3 \times 10^{-13}$) and type 2 diabetes ($p = 1.4 \times 10^{-6}$), and not for leiomyoma. It is therefore a generic property of complex-trait gene sets in these cells. The strongest endometriosis scDRS signal was in endometrial-type stromal states (eF2, eF3, dS4; $q = 0.02$), shared with leiomyoma. Their cell selection, labels and gene scoring differ from ours, and their code is public; this is a non-reproduction, not a refutation.

**Fallopian tube.**

To test a tubal origin of endometriosis-like epithelium we compared tubal secretory and ciliated epithelium [16] with endometrial epithelium of HECA (MAGMA gene-property, conditioned on mean expression in both atlases). Neither tubal secretory epithelium (KOLLER_EUR $p = 0.76$; 0.43 given endometrial epithelium) nor tubal ciliated epithelium (0.78; 0.25 given endometrial ciliated cells) was preferentially enriched, and no epithelial compartment of either organ reached $p < 0.05$ for any endometriosis GWAS. In a post hoc stromal comparison, tubal and uterine stroma were indistinguishable for KOLLER_EUR (0.037 and 0.046; difference $p = 0.46$) and attenuated each other when fitted jointly.

### Supplementary figures

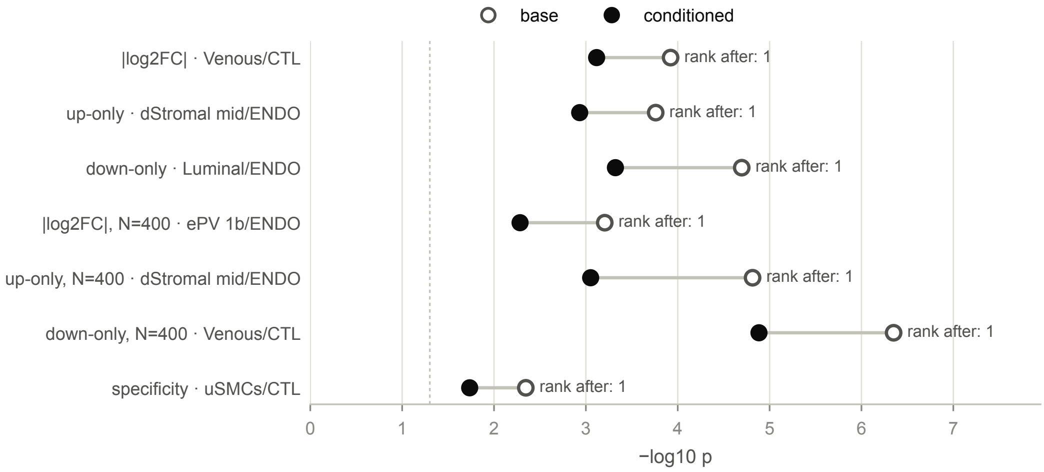


**Figure S1**: Conditioning on generic gene properties. First-ranked HECA label per family for KOLLER_EUR before and after conditioning on mean expression, expression breadth and gnomAD LOEUF (MAGMA gene-property analysis of binary set indicators); rank after conditioning on the right.

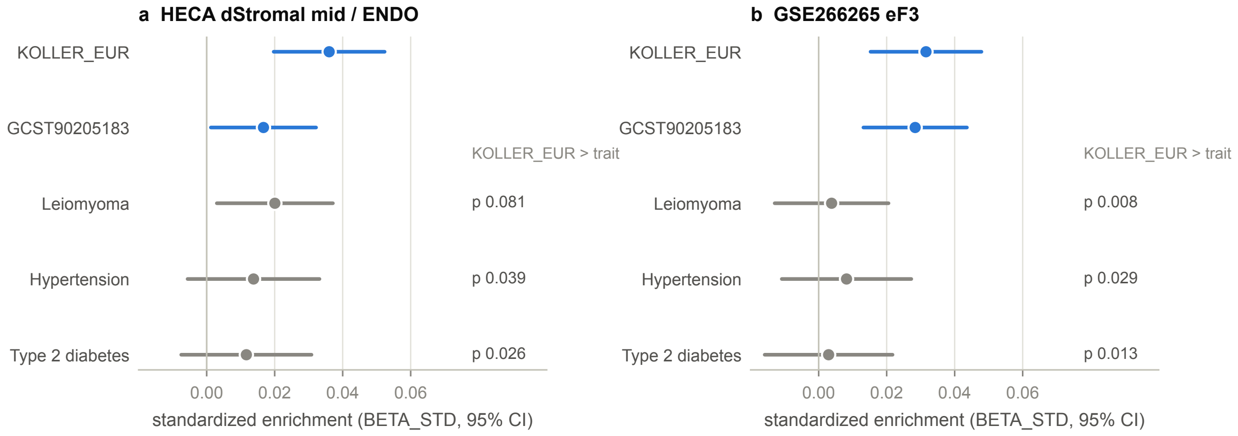


**Figure S2:** Trait contrasts with GWAS covariance. MAGMA enrichment of the size-matched up-only sets (N = 400) per GWAS; whiskers, 95% CI from the MAGMA standard error. Endometriosis GWAS in blue, control traits in gray. Right column, one-sided p that KOLLER_EUR exceeds the control trait, with the covariance of the two estimates taken from 1,000 paired genome rotations; the empirical rotation p was smaller in every case.

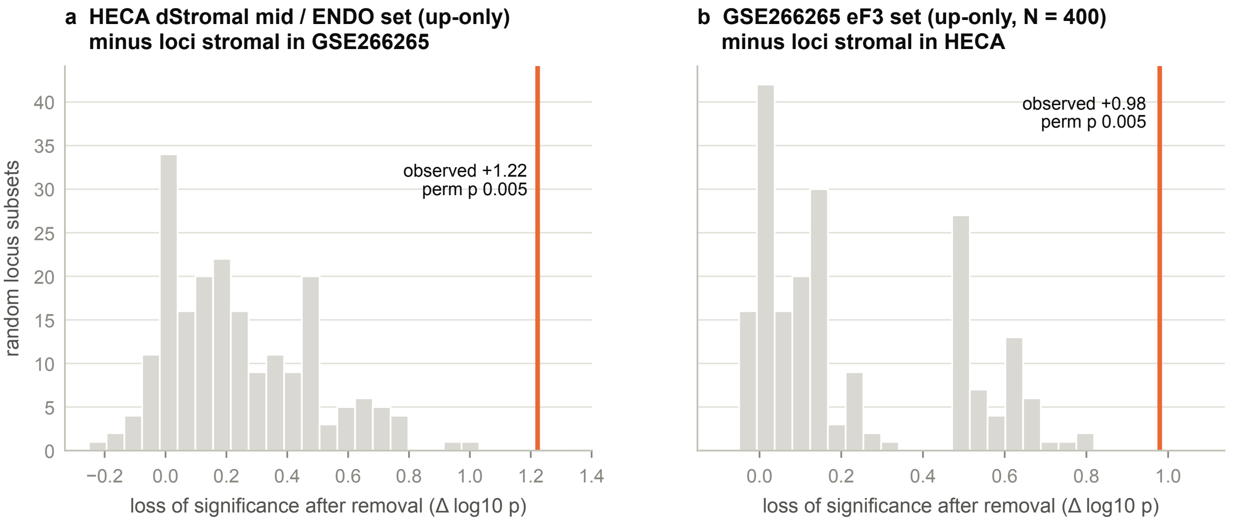


**Figure S3:** Cross-atlas locus test. KOLLER_EUR, MAGMA; genes within 500 kb of each removed lead SNP dropped from the set. (a) Loci classified as stromal with GSE266265 expression removed from the HECA up-only dStromal_mid/ENDO set. (b) Loci classified as stromal with HECA expression removed from the GSE266265 size-matched up-only eF3 set. Gray, 200 random equal-size subsets of the other loci; orange, observed.

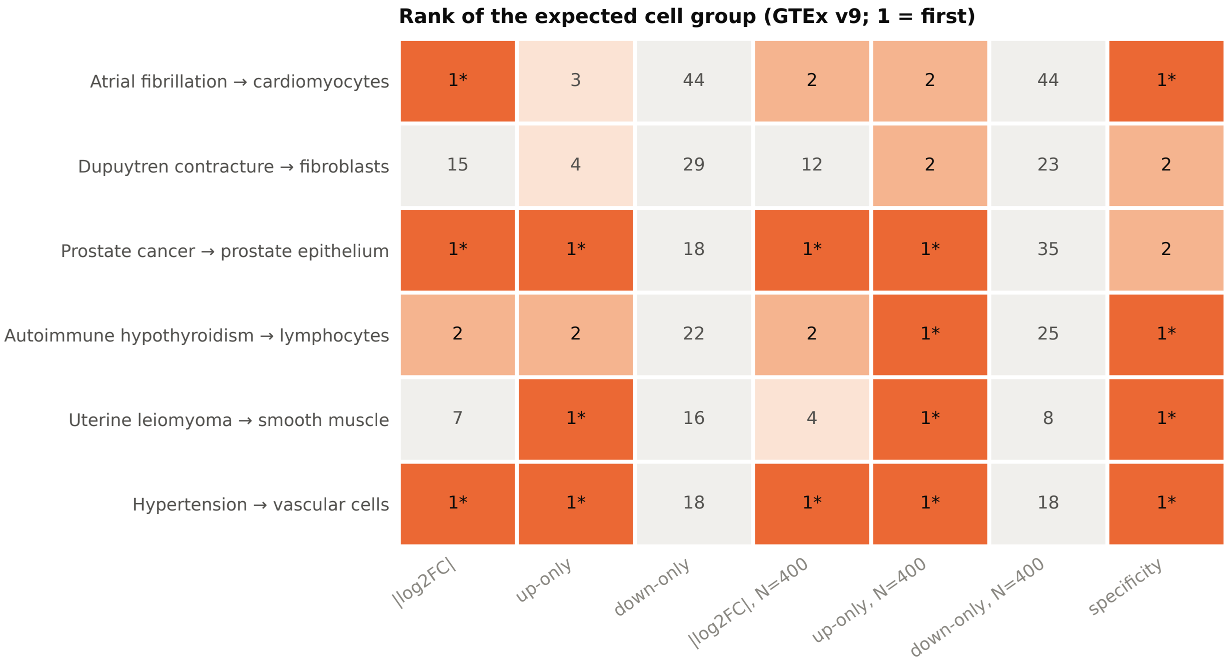


**Figure S4**: Known-source control panel. GTEx v9 snRNA-seq [24]; six FinnGen R13 GWAS, each with a cell group fixed before the run. Each cell gives the rank of the best label of the expected group under one marker family (MAGMA; asterisk, first with BH $q < 0.05$). Thresholded families use $|\log 2\ FC| > 1.2$ and adjusted $p < 0.01$ as in Koller et al.; $N = 400$ families are size-matched; specificity is the continuous share of expression. Label rules for prostate epithelium and lymphocytes as corrected after the label list was first printed; ranks under the registered rules differ only for size-matched up-only in prostate cancer (2) and the down-only families in autoimmune hypothyroidism.

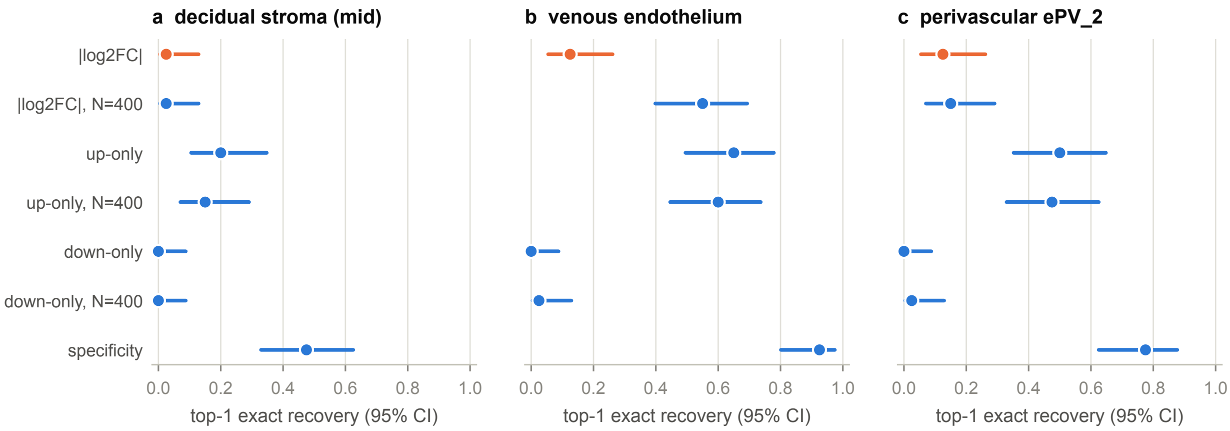


**Figure S5:** Benchmark intervals. Share of replicates (40 per truth) in which the planted state is ranked first, with Wilson 95% intervals, under specificity truths. The thresholded | log2 FC| rule of Koller et al. in orange.

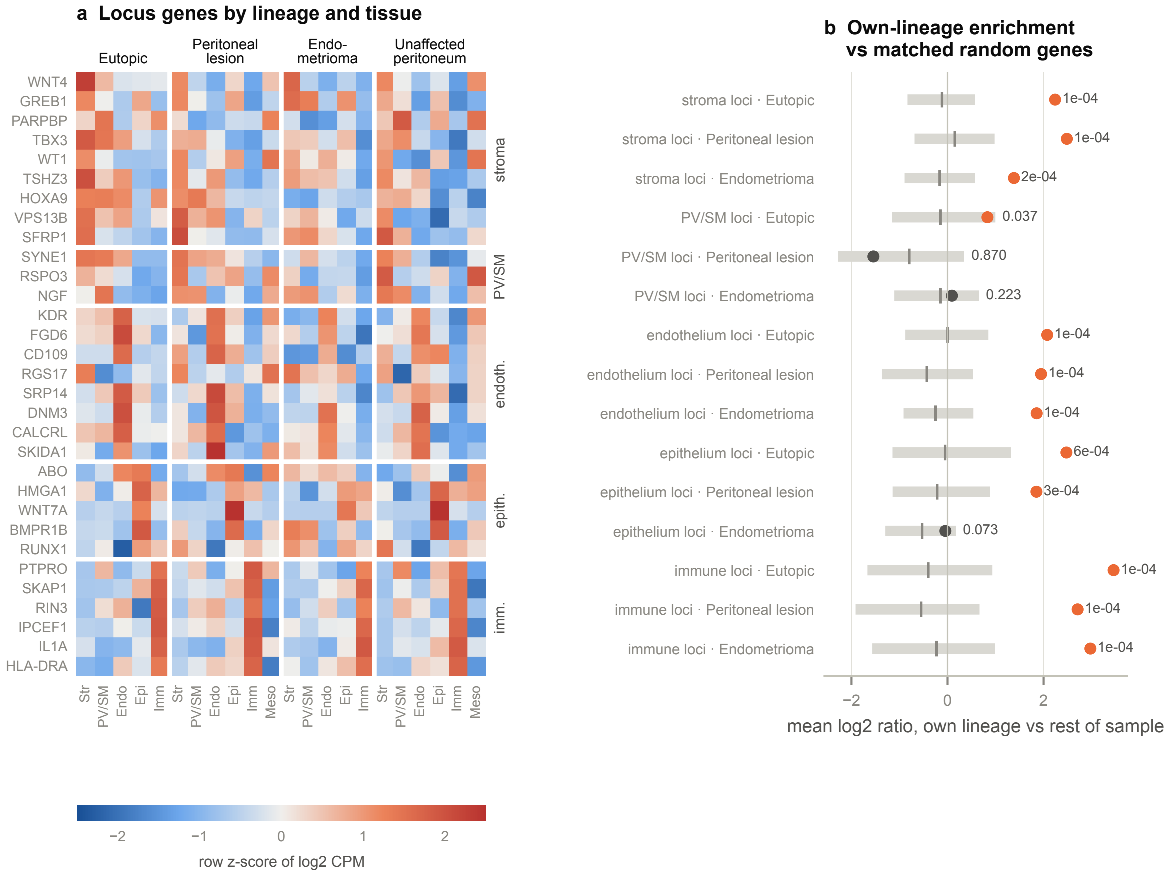


**Figure S6:** Genes at the endometriosis risk loci inside lesions. GSE213216 [13]; markerbased lineage pseudobulk (≥10 cells per sample and lineage), donor means. (a) Expression of each KOLLER_EUR locus gene, grouped by its HECA compartment, across lineages in eutopic endometrium (9 donors), peritoneal lesions (12), endometriomas (6) and unaffected peritoneum (2; shown for reference only); row z-score of log2 CPM. PARPBP and VPS13B are the expressed candidates of the IGF1 and OSR2 regions. (b) Mean log2 ratio of the locus genes in their own lineage versus the rest of the same sample (dot; orange, permutation $p < 0.05$), with the median (tick) and 95% range (bar) of 10,000 expression-matched random gene sets; one-sided permutation p at right.

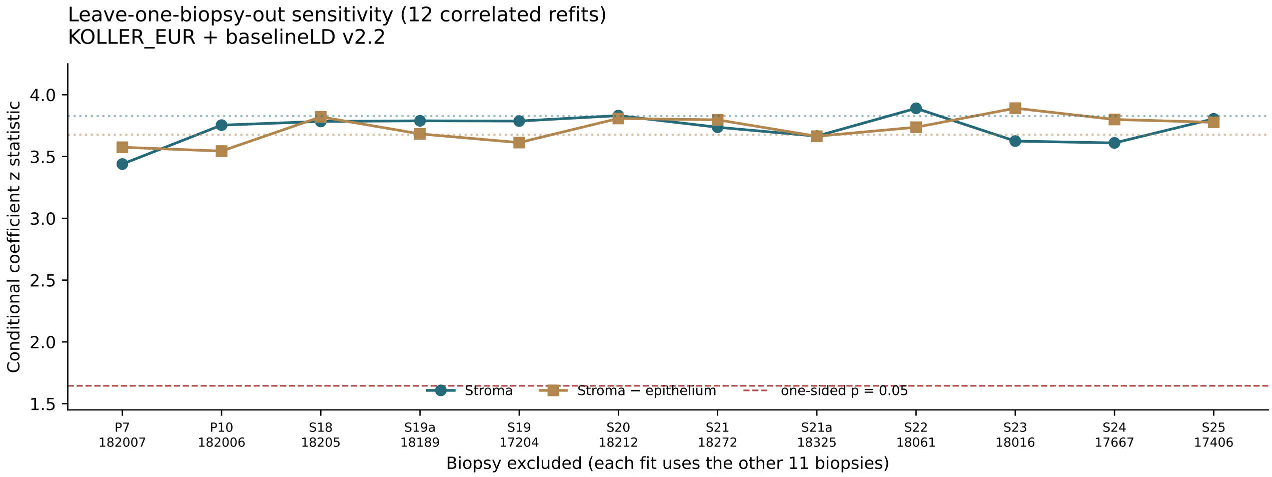


**Figure S7**: Post-result biopsy-exclusion robustness of the joint KOLLER_EUR chromatin model. Each point omits one of 12 distinct biopsy units (two technical runs were grouped for S19a_18189), then re-derives lineage-specific peaks, SNP masks, LD scores, and the joint S-LDSC fit. Full-data levels are indicated as a reference. Stromal and stroma-minus-epithelium coefficients remained positive and nominally significant in all 12 correlated refits. Cluster labels and the accessible-peak universe were held fixed; these estimates are not independent replications.

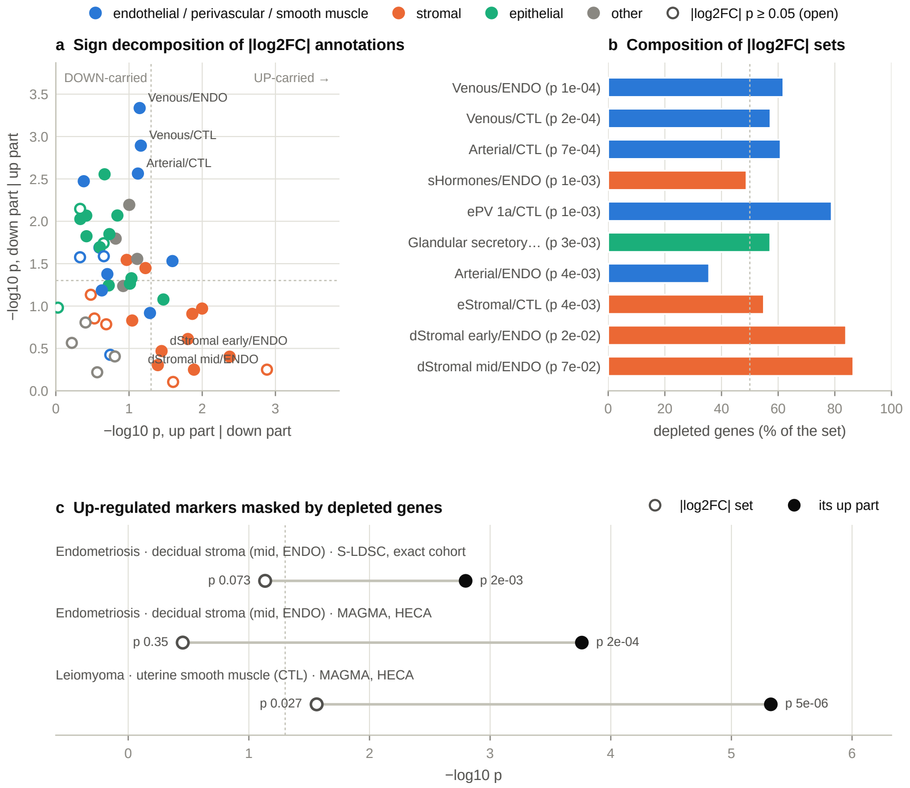


**Figure S8**: Depleted genes in | log2 FC| sets. Exact Koller cohort, KOLLER_EUR. (a) Joint S-LDSC of each cell type's up part and down part of its | log2 FC| set (dashed lines, p = 0.05; filled, | log2 FC| set nominally significant). (b) Share of depleted genes in the | log2 FC| sets of the top-ranked annotations and of decidual stroma (dashed line, 50%). (c) | log2 FC| set vs its up part for the expected source cell type of each trait: decidual stroma for endometriosis (S-LDSC exact cohort; MAGMA HECA) and uterine smooth muscle for leiomyoma (MAGMA HECA).

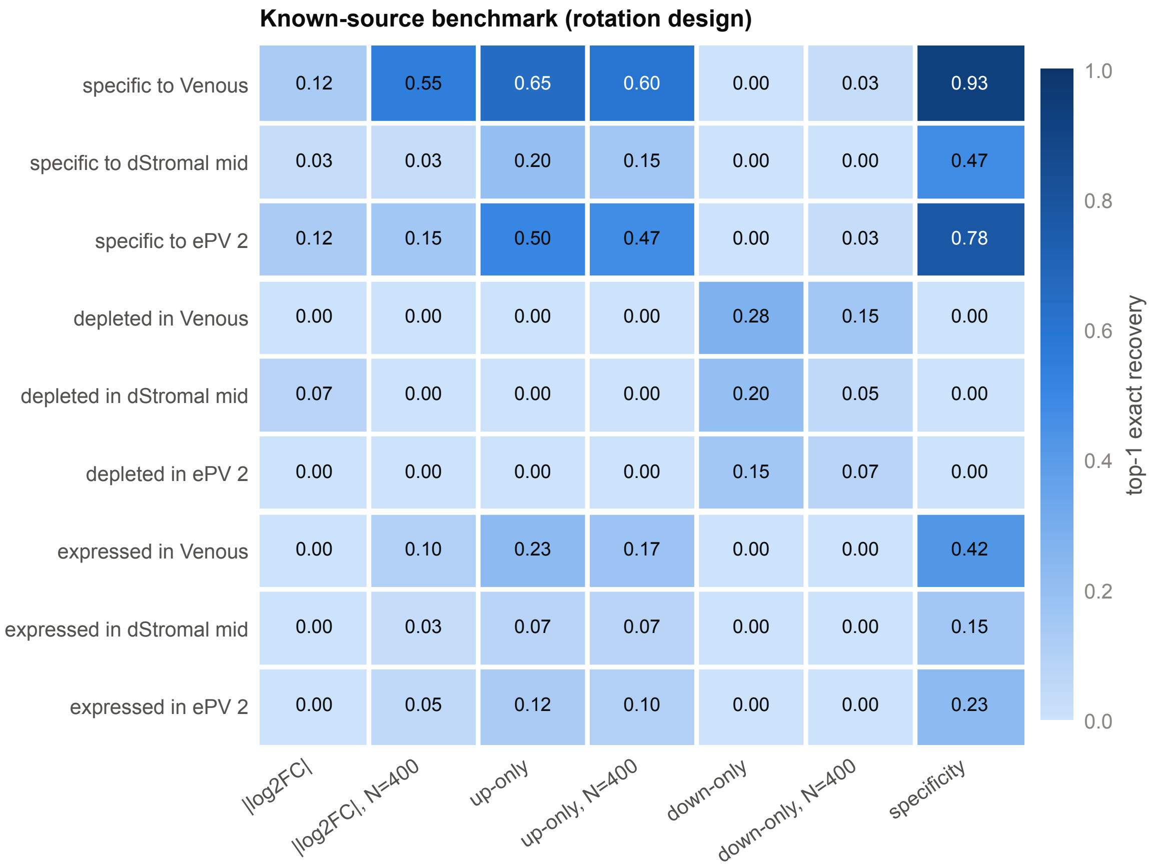


**Figure S9**: Known-source benchmark. Share of replicates in which the planted source state is ranked first (MAGMA, HECA, real KOLLER_EUR z-scores with a rotated annotation layer; 40 replicates per truth) for seven marker-selection families. Truths: genes specific to, depleted in, or highly expressed in the target state. The thresholded | log2 FC| rule is never the best family. Intervals are shown in Figure S5.

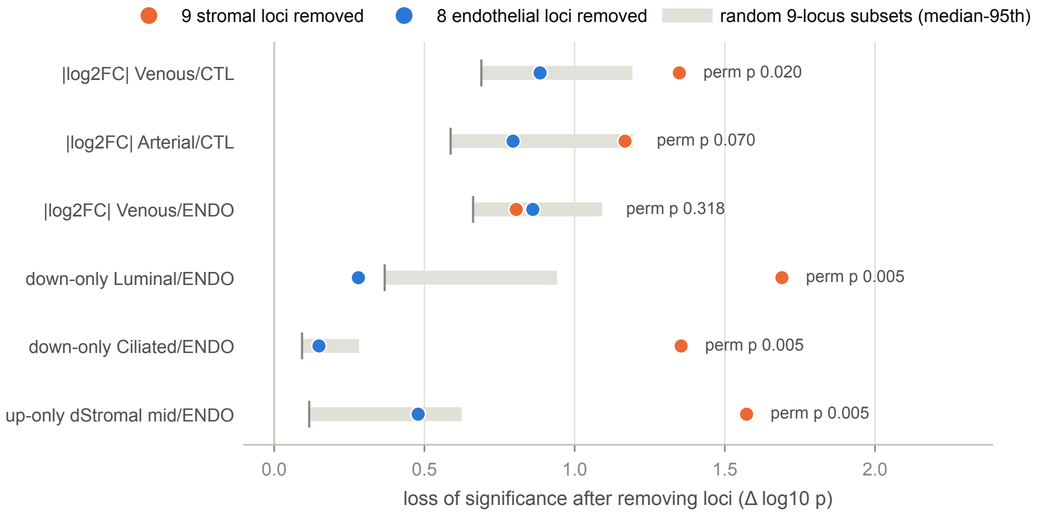


**Figure S10**: Nine stromal loci carry three different cell labels. KOLLER_EUR, MAGMA, HECA marker sets. Increase in log10 p of each first-ranked set after removing the genes within 500 kb of the nine stromal-assigned genome-wide loci or of the eight endothelial-assigned loci, against 200 random 9-locus subsets of the non-stromal loci (post hoc test).

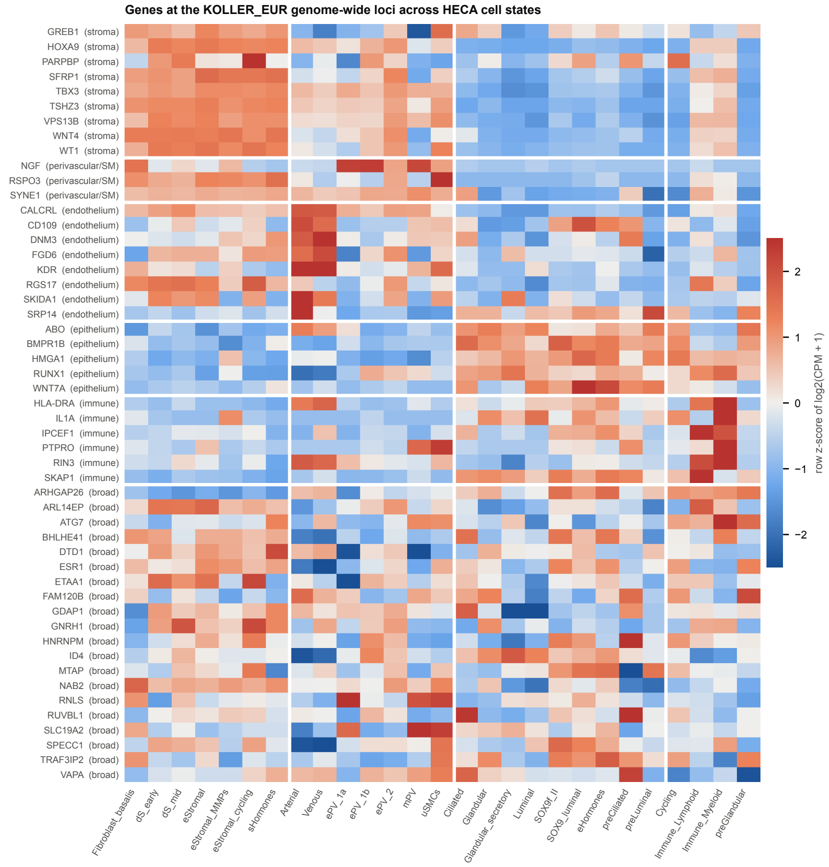


**Figure S11:** Expression of the locus genes across HECA cell states. Gene used for each KOLLER_EUR genome-wide locus (row z-score of log2(CPM + 1); control and endometriosis donors averaged), grouped by assigned compartment. Genes at the nine stromal loci peak in proliferative, hormone-responsive and decidual endometrial stroma.

### Supplementary tables

**Table S1:** Prospectively specified, sequentially registered follow-up, internal-audit, and original chromatin tests, with all unsupported outcomes retained. The original CH-1–CH-3 tests preceded inspection of their GWAS results; subsequent joint-model and leave-one-biopsy-out sensitivity analyses are reported separately and were not prospectively specified. Trait contrasts are covariance-aware.

| Group | Test | Prediction | Observed outcome | Verdict |
|---|---|---|---|---|
| Confirmatory | **H-EXACT** | Count-matched cohort: ABS first endothelial, UP first decidual; leiomyoma smooth muscle first only with UP | Venous/ENDO q 0.01; dStromal_mid/ENDO first, q 0.08; smooth muscle ranks 1 vs 8 | Supported (rank-based; nominal) |
| Confirmatory | **H-ABS** | All endothelial ABS sets are >50% depleted | Arterial/ENDO 36% in exact cohort; all >50% in sensitivity cohort | Not supported |
| Confirmatory | **H-SIGN** | All endothelial hits down-carried and decidual hits up-carried | 3/4 in each group | Not supported |
| Confirmatory | **K8 (G4 prediction)** | UP dStromal_mid not separable from perivascular ePV_2 | Separable; survives 6/6 top sets and 11/11 vascular sets | Not supported (stroma separable) |
| Confirmatory | **Benchmark calibration** | FWER ≤5% for all seven families | 0.3–1.7%; all upper bounds <5% | Supported |
| Confirmatory | **Leiomyoma control (HECA, GTEx)** | Smooth muscle first with up-type rules | Yes; ABS ranks 2 and 7 | Supported |
| Confirmatory | **L2c** | Stromal loci carry the UP decidual signal | Permutation p=0.020 | Supported |
| Follow-up | **H-STRVEN** | Stromal loci carry the ABS venous hit more than chance and more than endothelial loci | p=0.020; +1.35 vs +0.89 | Supported |
| Follow-up | **H-NONSPEC** | Down-type families more shared across traits | HECA −0.09; GTEx +0.20 (CI reaches −0.00) | Not supported |
| Follow-up | **H-ABSDOWN** | ABS sets mostly depleted and rankings follow down-sets | GTEx 33% depleted | Not supported |
| Follow-up | **H-GENERIC** | Gene properties remove more down-type than up-type significance | Holds in HECA; reversed in GTEx | Not supported |
| Follow-up | **H-DSTROMA** | UP decidual remains nominally significant after generic gene-property conditioning | $p=1.2\times10^{-3}$ and $8.9\times10^{-4}$ | Supported |
| Follow-up | **H-K7REP** | Top ABS hits mostly down-carried in MAGMA | 0.33 GTEx; 0.37 HECA | Not supported |
| Follow-up | **H-MASK** | Leiomyoma sets above smooth muscle are down-carried | HECA yes; GTEx 2/6 | Not supported |
| Follow-up | **H-ROBUST-FLIP** | UP first decidual in ≥6/8 threshold settings | 5/8 decidual; 8/8 stromal | Not supported |
| Follow-up | **H-ROBUST-CTRL** | Leiomyoma smooth muscle first with UP in ≥6/8 and more often than ABS | 8 vs 2 | Supported |

Continued on next page

Table S1 (continued)

| Group | Test | Prediction | Observed outcome | Verdict |
|---|---|---|---|---|
| Follow-up | **H-DS-SPECIFIC** | Decidual (UPN) stronger for endometriosis than T2D and hypertension | p=0.026, 0.039 | Supported |
| Follow-up | **H-REPL-FLIP** | GSE266265 reproduces rule-dependent first-ranked cell | UP first vascular smooth muscle; ABS did not give expected first-ranked alternative | Not supported |
| Follow-up | **H-REPL-CTRL** | GSE266265 hypertension vascular first with up-type markers | Yes | Supported |
| Follow-up | **H-ORIGIN** | GSE266265 stroma above perivascular, mesothelial-like, epithelial | Ranks 1, 2, 5, 11 | Supported |
| Follow-up | **H-ORIGIN-SPECIFIC** | eF3 stronger for endometriosis than T2D and hypertension | p=0.013, 0.029 | Supported |
| Follow-up | **H-STROMA-SEP** | eF3 survives conditioning on each perivascular/mesothelial set | $p=1–3\times10^{-4}$ | Supported |
| Follow-up | **H-EF3-ROBUST** | eF3 survives gene-property conditioning | $p=2.5\times10^{-3}$ | Supported |
| Follow-up | **H-MESO** | Endometrial-type stroma above real mesothelium and peritoneal fibroblasts | Ranks 1, 3, 6; all p≥0.05 | Rank met; nominal |
| Follow-up | **H-LESION-LOC** | GSE213216 stromal-locus genes enriched in lesion stroma | $p=1\times10^{-4}$, $2\times10^{-4}$ | Supported |
| Follow-up | **H-PANEL** | Six known-source traits: UP first in ≥4 and more often than ABS | 3 vs 3 | Not supported |
| Follow-up | **H-PANEL-N** | Size-matched UP first more often than size-matched ABS | 4 vs 2 | Supported |
| Follow-up | **H-LESION-REP** | GSE179640 stromal-locus genes enriched within lesions | $p=1\times10^{-4}$ in both lesion types | Supported |
| Follow-up | **H-HOST** | GSE179640 lesion stroma above adjacent-peritoneum stroma | Difference +0.01; p=0.26 | Not supported |
| Follow-up | **H-MESO-REP** | GSE179640 endometrial stroma above mesothelium | Mesothelium first, p=0.018 | Not supported |
| Follow-up | **H-NE-EAS** | East Asian GWAS: eF3 enriched with size-matched UP | p=0.076 | Not supported |
| Follow-up | **H-NE-ORIGIN** | East Asian GWAS: stroma above alternative compartments | eF1 first, p=0.01 | Supported (nominal) |
| Audit | **H-COV-CONTRAST** | HECA decidual and GSE266265 eF3 stronger for endometriosis than T2D and hypertension in covariance-aware and rotation tests | HECA p=0.026, 0.039; eF3 p=0.013, 0.029 | Supported (nominal) |
| Audit | **H-LOCUS-X** | Loci classified as stromal in one atlas carry stromal set of the other | Permutation p=0.005 in both directions; κ=0.77 | Supported |
| Audit | **H-LOO-GSE** | GSE266265 stroma first in ≥90% of 60 donor exclusions for KOLLER and GCST | 48/60 and 60/60; eF3 $p\leq10^{-3}$ in all runs | Not supported |
| Audit | **H-LOO-HECA** | HECA status-split: decidual first in all but one of 7 donor exclusions | 6/7 decidual; stroma 7/7; states single-donor dominated | Supported (rule) |

Continued on next page

Table S1 (continued)

| Group | Test | Prediction | Observed outcome | Verdict |
|---|---|---|---|---|
| Audit | **H-HECA-PB** | HECA status-split donor-paired pseudobulk ranks stroma first | Perivascular ePV_1b first, p=0.02 | Not supported |
| Audit | **H-HECA-BAL** | HECA pooled donor-balanced: stroma first in at least 13/14 exclusions | 14/14 | Supported |
| Audit | **H-HECA-PB2** | HECA pooled donor-paired pseudobulk: stroma first in at least 13/14 | 12/14 | Not supported |
| Chromatin | **CH-0 (pre-GWAS audit)** | GSE183771 annotations pass composition, lift-over, biopsy counts and culture-overlap gates | Passed after two disclosed labeling revisions; stromal culture overlap 67% vs 32% | Passed |
| Chromatin | **CH-1** | KOLLER_EUR stroma-specific accessible chromatin enriched | z=4.11, $p=2\times10^{-5}$; GCST 4.27, FinnGen 2.51; HTN/T2D null | Supported |
| Chromatin | **CH-2** | Stroma above epithelium in a single model | z=3.72; $p=1\times10^{-4}$ | Supported |
| Chromatin | **CH-3** | Stroma first among four lineage annotations | z: stroma 4.11; endothelium 0.15; immune −1.11; epithelium −2.49 | Supported |
| Descriptive | **K4 ABS joint** | Decidual sets conditioned on leading non-focus sets | 0/7 survive; non-focus sets survive all | Descriptive |
| Descriptive | **Donor composition** | Largest donor share by status-specific HECA label | >80% in 16/45 groups; median 2.3 effective donors | Descriptive |
| Descriptive | **LDSC rg** | GWAS overlap from cross-trait intercepts | KOLLER vs GCST 0.53; vs FinnGen 0.69; GCST vs FinnGen 0.05 | Descriptive |
| Expanded, descriptive | **Nine-phenotype leader metric** | Change in first-ranked annotation between UP and ABS across the nine fixed FinnGen traits | 4/9 annotations; 2/9 broad compartments; the heart-failure shift lacks BH-supported leaders | Descriptive (not accuracy) |
| Expanded, exploratory | **Six proxy ranks** | Two-sided exact sign-flip of signed log ranks for six predeclared broad cell-type proxies | Four UP improvements, one worsening, one tie; five nonzero; $p = 0.125$ | Inconclusive |
| Expanded, exploratory | **Five rules vs ABS** | Five two-sided sign-flip comparisons (DOWN, ABSN, UPN, DOWNN, SPEC), with family-wise Holm adjustment | No adjusted $p < 0.05$; best unadjusted UPN $p = 0.0625$ | Inconclusive |

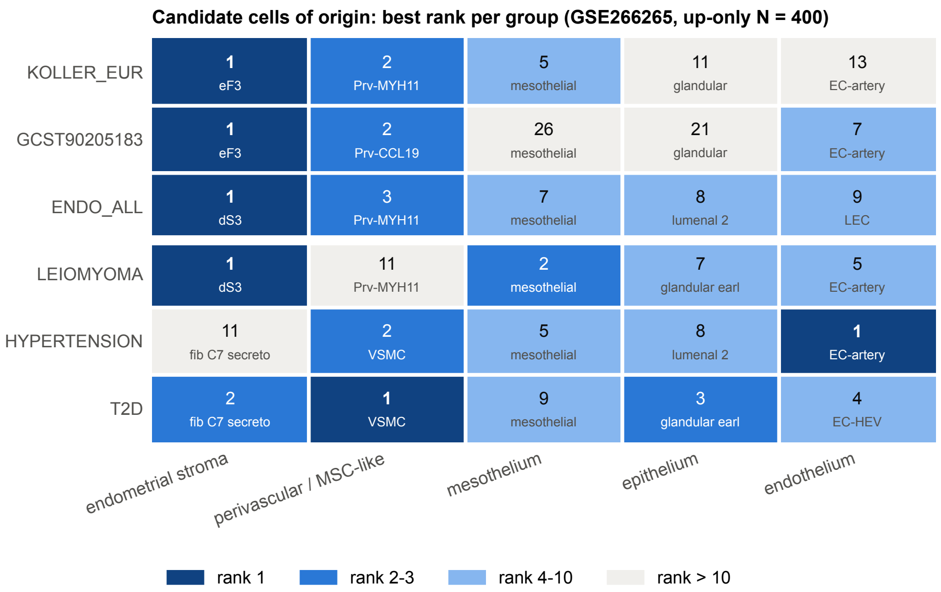


**Figure S12**: Candidate cells of origin in an independent atlas. Best rank of each candidate-origin group per GWAS in GSE266265 (60 donors; donor-paired pseudobulk; size-matched up-only sets, N = 400; MAGMA). Endometrial stroma (eF3; dS3 for FinnGen) ranks first for all three endometriosis GWAS; perivascular cells are second. The "mesothelial" label of this atlas is not mesothelium by marker expression and is not interpreted.

**Table S2**: Nine new FinnGen R13 phenotypes: first-ranked GTEx v9 cell-type annotations under absolute-fold-change (ABS) and up-regulated (UP) marker selection. Each $q$ is adjusted within its 45-cell marker family; the rank correlation $\rho$ is Spearman across those 45 annotations.

| Phenotype | ABS leader | $q_{ABS}$ | UP leader | $q_{UP}$ | $\rho$ |
|---|---|---|---|---|---|
| Psoriasis | myeloid dendritic cell | 0.024 | myeloid dendritic cell | 0.002 | 0.837 |
| COPD | myeloid dendritic cell | 0.005 | myeloid dendritic cell | 0.005 | 0.624 |
| Asthma | myeloid dendritic cell | 0.041 | T cell | 0.047 | 0.728 |
| Breast cancer | fibroblast of cardiac tissue | 0.003 | bronchial epithelial cell | 0.021 | 0.737 |
| Coronary disease | endothelial cell of vascular tree | 0.00019 | endothelial cell of vascular tree | 0.003 | 0.567 |
| Non-small-cell lung cancer | luminal cell of prostate epithelium | 0.333 | luminal cell of prostate epithelium | 0.148 | 0.839 |
| Atopic dermatitis | myeloid dendritic cell | 1e-05 | myeloid dendritic cell | 1.7e-06 | 0.843 |
| Heart failure | slow muscle cell | 0.071 | macrophage | 0.161 | 0.752 |
| Varicose veins | cardiac endothelial cell | 1.3e-06 | endothelial cell of lymphatic vessel | 1.1e-05 | 0.523 |

**Table S3**: Six broad cell-type proxy labels defined in an analysis-plan addendum before calculation of the nine new genetic cell-rank results. These are biological proxies, not uniquely causal cell identities. Lower cell rank indicates higher priority; signed effect is $\log(\text{rank}_{UP}) - \log(\text{rank}_{ABS})$. The secondary two-sided exact sign-flip test over five nonzero pairs gave $p = 0.125$.

| Phenotype | Predeclared GTEx cell-type proxy | ABS rank | UP rank | Log-rank difference |
|---|---|---|---|---|
| Psoriasis | basal cell of epidermis | 11 | 12 | +0.087 |
| COPD | pulmonary alveolar epithelial cell | 27 | 21 | -0.251 |
| Breast cancer | luminal epithelial cell of mammary gland | 23 | 21 | -0.091 |
| Atopic dermatitis | epidermal cell | 25 | 19 | -0.274 |
| Heart failure | cardiac muscle cell | 33 | 30 | -0.095 |
| Varicose veins | endothelial cell of vascular tree | 2 | 2 | +0.000 |

**Table S4**: Post-result full joint scATAC S-LDSC coefficients with baselineLD v2.2. Models include the ALL-peak control and four lineage-specific top-10% annotations simultaneously; $p_+$ is a one-sided nominal test. The GWAS share participants, and leiomyoma shows that stromal accessibility is not endometriosis-specific.

| GWAS | Stromal $z$ | $p_+$ | Stroma−epithelium $z$ | $p_+$ |
|---|---|---|---|---|
| KOLLER EUR | 3.828 | 6.5e−5 | 3.677 | 0.00012 |
| GCST90205183 | 4.055 | 2.5e−5 | 3.317 | 0.00046 |
| FinnGen endometriosis | 1.979 | 0.024 | 2.659 | 0.0039 |
| Leiomyoma | 2.418 | 0.0078 | 2.982 | 0.0014 |
| Hypertension | 0.177 | 0.43 | -0.768 | 0.78 |
| Type 2 diabetes | -0.797 | 0.79 | -1.687 | 0.95 |

**Table S5**: Full joint scATAC S-LDSC sensitivity across two genomic backgrounds. All five ATAC annotations are in each joint model; $p_+$ denotes an exploratory one-sided nominal test. The genetic samples overlap.

| Background | Trait | Stromal $z$ | $p_+$ | Stromal–epithelial $z$ | $p_+$ |
|---|---|---|---|---|---|
| baseline 1.2 | KOLLER EUR | 3.891 | 5e−5 | 3.781 | 7.8e−5 |
| baseline 1.2 | GCST90205183 | 4.221 | 1.2e−5 | 3.456 | 0.00027 |
| baseline 1.2 | FinnGen endometriosis | 2.163 | 0.015 | 2.748 | 0.003 |
| baseline 1.2 | Leiomyoma | 2.286 | 0.011 | 2.956 | 0.0016 |
| baseline 1.2 | Hypertension | 0.217 | 0.41 | -0.476 | 0.68 |
| baseline 1.2 | Type 2 diabetes | -0.789 | 0.79 | -1.400 | 0.92 |
| baselineLD 2.2 | KOLLER EUR | 3.828 | 6.5e−5 | 3.677 | 0.00012 |
| baselineLD 2.2 | GCST90205183 | 4.055 | 2.5e−5 | 3.317 | 0.00046 |
| baselineLD 2.2 | FinnGen endometriosis | 1.979 | 0.024 | 2.659 | 0.0039 |
| baselineLD 2.2 | Leiomyoma | 2.418 | 0.0078 | 2.982 | 0.0014 |
| baselineLD 2.2 | Hypertension | 0.177 | 0.43 | -0.768 | 0.78 |
| baselineLD 2.2 | Type 2 diabetes | -0.797 | 0.79 | -1.687 | 0.95 |

**Table S6**: Post-result leave-one-biopsy-out stability of the KOLLER_EUR joint scATAC model with baselineLD v2.2. Peaks and LD scores were recomputed after each exclusion; fixed cell labels and fixed accessible-peak universe. All tests are nominal and dependent.

| Excluded biopsy | Stromal $z$ | $p_+$ | Stroma–epithelium $z$ | $p_+$ |
|---|---|---|---|---|
| P7_182007 | 3.439 | 0.00029 | 3.575 | 0.00017 |
| P10_182006 | 3.754 | 8.7e−5 | 3.544 | 0.0002 |
| S18_18205 | 3.784 | 7.7e−5 | 3.821 | 6.7e−5 |
| S19a_18189 | 3.789 | 7.6e−5 | 3.683 | 0.00012 |
| S19_17204 | 3.787 | 7.6e−5 | 3.613 | 0.00015 |
| S20_18212 | 3.831 | 6.4e−5 | 3.808 | 7e−5 |
| S21_18272 | 3.737 | 9.3e−5 | 3.797 | 7.3e−5 |
| S21a_18325 | 3.665 | 0.00012 | 3.664 | 0.00012 |
| S22_18061 | 3.890 | 5e−5 | 3.737 | 9.3e−5 |
| S23_18016 | 3.625 | 0.00014 | 3.891 | 5e−5 |
| S24_17667 | 3.610 | 0.00015 | 3.800 | 7.2e−5 |
| S25_17406 | 3.805 | 7.1e−5 | 3.777 | 7.9e−5 |
